# CD4+ tissue-resident memory Th17 cells are a major source of IL-17A in Spondyloarthritis synovial tissue

**DOI:** 10.1101/2024.10.11.617865

**Authors:** Feng Liu, Hui Shi, Jason D Turner, Rachel Anscombe, Jiaqi Li, Takuya Sekine, Ariane Hammitzsch, Devika Agarwal, Christopher Mahony, Jiewei Chen, Ben Kendrick, Dajiang Du, Qiang Tong, Lihua Duan, Kyla Dooley, Hai Fang, Ilya Korsunsky, Roopa Madhu, Adam P Cribbs, Cartography consortium, Matthias Friedrich, Brian D. Marsden, Yi-Ling Chen, Graham Ogg, Anna Adams, Warner Chen, Steven Leonardo, Fiona E. McCann, Christopher D Buckley, Terence Rooney, Thomas Freeman, Holm H. Uhlig, Calliope Dendrou, Adam Croft, Andrew Filer, Paul Bowness, Liye Chen

**Author notes:** Equal contributions (Feng Liu, Hui Shi).

## Abstract

**Objectives:** Interleukin (IL)-17A is a key driver of Spondyloarthritis (SpA) joint pathology. We aimed to identify its cellular source in synovial tissue from patients with two forms of SpA namely Axial SpA (AxSpA) and Psoriatic arthritis (PsA).

**Methods:** Synovial tissue from patients with SpA was profiled using single-cell RNA sequencing (scRNA-seq: AxSpA, n=5 and PsA, n=6) or spatial RNA profiling (PsA, n=4). CellPhoneDB was used to infer cell–cell communication. Tissue resident memory Th17 (TRM17)-like cells were generated *in vitro* using blood memory CD4+ T cells from SpA patients. An epigenetic inhibitor library, siRNA and clustered regularly interspaced short palindromic repeats (CRISPR) were used to identify epigenetic regulator(s) for TRM17.

**Results:** scRNA-seq showed that *CD4+CXCR6+* TRM17 cells are the predominant spontaneous *IL17A* producers in SpA synovium. Cell-cell communication and single-cell spatial analysis support the interaction between TRM17 and *CLEC10A+* dendritic cells, which were activated in SpA. Both sublining and lining fibroblasts in SpA synovium showed evidence of IL-17A activation. In vitro-generated CD4+ TRM17-like cells phenocopied joint tissue TRM17, producing IL-17A/F upon T cell receptor (TCR) stimulation, which was further enhanced by cytokines. Perturbation of BRD1 inhibited the generation of TRM17-like cells.

**Conclusions:** CD4+ TRM17 cells are the predominant source of IL-17A in SpA synovial tissue. TCR stimulation is essential for the secretion of IL-17A by CD4+TRM17-like cells. The epigenetic regulator BRD1 contributes to the generation of CD4+TRM17. Depleting CD4+TRM17 cells in SpA is thus a therapeutic strategy with potential to induce long-term remission.

**Key messages:** *What is already known on this topic:* Interleukin (IL)-17A plays a key role in the immunopathogenesis of Spondyloarthritis (SpA), but its cellular source in joint tissue has not been determined previously. The induction and accumulation of CD4+ tissue resident memory Th17 (TRM17) cells following the clearance of pathogens has been described in skin, lung and kidney. Whether CD4+ TRM17 cells also accumulate in the joint and contribute to the pathology of SpA is not clear.

*What this study adds:*

I. CD4+ TRM17 cells are present in SpA synovial tissue and are the predominant source of *IL17A*
II. CD4+ TRM17 cells in SpA joints express *IL17A* without any *in vitro* exogenous stimulation
III. T cell receptor (TCR) rather than cytokine stimulation is essential for IL-17A production by CD4+ TRM17-like cells
IV. The epigenetic regulator BRD1 contributes to the generation of CD4+ TRM17-like cells.

*How this study might affect research, practice or policy:* Our findings identify CD4+ TRM17 cells as the primary source of IL-17A in SpA synovium, a previously unrecognized role for these cells. Key questions remain: How do CD4+ TRM17 cells relate to IL-17A producers in synovial fluid? What mechanisms induce and maintain them in the joint? How do they interact with other cells to promote arthritis? These questions warrant further investigation. In addition, our data suggest that targeting CD4+ TRM17 cells, the “factory” of IL-17A in SpA synovial tissue, has the potential to induce long-term remission, encouraging future efforts to develop new therapies to deplete CD4+ TRM17 cells in SpA.

## INTRODUCTION

The spondyloarthritides (SpA) are a group of common inflammatory arthritides that include Axial SpA (AxSpA) and psoriatic arthritis (PsA). The SpA share pathological drivers including interleukin 17 (IL-17) and tumor necrosis factor (TNF). Drugs targeting IL-17 are effective in over 50% of SpA patients but require continued long-term administration, and therapies to target the cells producing IL17 have not yet been developed.

IL-17A is known to promote tissue inflammation, at least in part through fibroblast activation. Among the different forms of IL-17 (A-F), IL-17A is the key driver in joint inflammation whereas a major role for IL-17F in skin inflammation has been supported by recent trials[1]. IL-17A can be produced by a variety of immune cells, including conventional T cells, innate-like lymphocytes and innate lymphoid cells (ILCs). Whether a specific cell type dominates IL-17A production in SpA is debated[2]. Different studies have implicated all three subsets of innate-like lymphocytes (γδ T cells, iNKT cells, and MAIT cells) and CD8+ tissue-resident memory T cells as the sources of IL-17A in SpA joints[3–10]. Increasing evidence, however, is highlighting a role for CD4+ tissue-resident memory Th17 (TRM17) cells as key IL-17A producers in tissues such as the skin, lung and kidney[11–13]. Furthermore, CD4+ Th17 cells expressing CXCR6, an established marker for tissue residency, have been linked to brain inflammation in a murine model of multiple sclerosis [14]. It is therefore possible that a role for CD4+ TRM17 cells in SpA may have been missed in previous studies focused on synovial fluid samples and/or utilising potent *in vitro* stimulation to determine the source of IL-17.

Notably CD4+CXCR6+ Th17 cells in inflamed brain tissue have been found to exhibit a distinct epigenetic profile that is different from homeostatic Th17 cells with stem-like features[14]. Considering the established role of epigenetic modifiers in the regulation of T cell differentiation and function[15,16], it is plausible to predict that epigenetic regulators could be involved in the generation of CD4+ TRM17 cells.

Through profiling un-stimulated synovial tissue using single-cell RNA sequencing and spatial RNA profiling, we here show that CD4+ CXCR6+ TRM17 cells are the predominant source of IL-17A in SpA synovial tissue. We also establish a model to generate CD4+ TRM17-like cells *in vitro* and show that T cell receptor (TCR) engagement, rather than cytokines, is essential for these cells to produce IL-17A and IL-17F. Lastly, we show that the epigenetic regulator bromodomain-containing protein 1 (BRD1) contributes to the generation of CD4+ TRM17-like cells.

## METHODS

### Patient Recruitment

Fresh synovial tissue was obtained at arthroplasty or synovial biopsy was collected from patients attending the Oxford University Hospitals (OUH), Queen Elizabeth Hospital and Shanghai Sixth People’s Hospital with appropriate consent and ethical approvals. All patients included in this study met the criteria of the Assessment of Spondyloarthritis International Society (ASAS) for AxSpA or CASPAR criteria for PsA. Patients currently or previously receiving IL17i therapy were excluded from the study population. The demographics of AxSpA and PsA patients recruited for this study are shown in **Table S1**. **Computational analysis of scRNA-seq data** The paired reads obtained were mapped to the hg38 reference genome to generate gene expression matrices using CellRanger. For PsA samples (from the Cartography Consortium), the full data set was analysed in Panpipes [17] and T cell compartment was partitioned out for downstream analyses along with the AxSpA data. T cells of PsA samples and full dataset of AxSpA samples were then merged and analyzed using the Seurat R package (v4.0.5). Cells with low-quality profiles were excluded based on the number of detected genes, the percentage of mitochondrial RNA among total UMIs, and the total number of UMIs. The raw read counts were normalized using the NormalizeData function, and variable genes were identified for each sample. To identify cell types, we mapped our cells into a synovium reference map built by Fan Zhang et. al.[18] using Symphony algorithm. For the reclustering analysis of cellular subsets, a certain population were extracted and data from different sample were subject to FastMNN algorithms for integration. We then used FindNeighbors and FindCluster functions to cluster cells based on global transcriptional profile. For differential expression analysis, the FindMarkers function was used to test the normalized data with the default Wald test method. Unless specified, default parameters were used for each function.

For cell-cell interaction inference, normalised expression matrix and cluster annotations were exported from the Seurat object as input for cellphonedb. Cellphonedb v5.0.0 was used as a ligand-receptor interaction database. ‘statistical analysis’ method from Cellphonedb tools was used to assess the strength and significance of interactions.

### CosMx spatial analysis

Synovial biopsy tissue sections from eight ultrasound guided biopsies were distributed across two flow cells and run on the CosMx Spatial Molecular Imager. Cell segmentation and initial quality control were completed in the AtoMx Spatial Informatics Platform before exporting and using R (v4.4.0) and Seurat (v5.1.0) for further quality control, clustering, and cluster annotation. Data were processed through iterations of clustering, annotation, identification and removal of multiplets/poorly segmented cells based on gene expression profiles; number of genes detected; and number of transcripts detected, and subclustering until low quality/poorly segmented cells were minimised in the dataset.

### Patient and Public Involvement

We have actively engaged the patients with SpA through the Oxford Patient Engagement Network for Arthritis and Musculoskeletal Conditions (OPEN ARMS, https://www.ndorms.ox.ac.uk/get-involved/open-arms-1/open-arms) and the Botnar AxSpA day. We will discuss with patients about the impact of findings from this study and co-develop a dissemination plan to maximize the potential benefits of our findings to patients.

### Additional methods

Information for Th17 expansion assay for the screen of epigenetic inhibitor library, siRNA screen of the targets of OF-1, intracellular cytokine staining (ICS) and enzyme-linked immunosorbent assays (ELISA) are available in **supplementary methods**.

## RESULTS

### Tissue resident memory CD4+ T cells are the main source of *IL17A* in SpA joint tissue

To investigate the cellular origin of IL-17A in the inflamed joints of SpA patients, we performed single-cell transcriptomic profiling on synovial tissue from 5 AxSpA and 6 PsA patients (demographics shown in Table S1A and B). After removing low-quality cells, we obtained 85,307 cells (Figure S1). These cells were categorized into T cells, natural killer (NK), mast and innate lymphoid cells (ILCs), B and plasma cells, myeloid cells, stromal cells, and endothelial cells (Figure 1A). Within this cellular milieu we found almost all *IL17A*+ cells are localized within the *CD4+* T cell subset (Figure 1B). Notably, the vast majority of the *IL17A*+ cells did not express *CD8A* (encoding the CD8 protein), *TRAV1-2* (encoding the T cell receptor α-variable 1-2 protein, abundant in mucosal-associated invariant T [MAIT] cells), or *TRDC* (encoding the delta constant region of the T cell receptor, characteristic of γδ T cells) (Figure 1C). As ILCs especially ILC3 cells are known to be capable of producing IL-17A, we investigated further into ILCs via the re-clustering of the cluster that contains NK, mast and ILCs (Figure S2A). Albeit expressing high levels of *RORC* and *IL23R, IL7R+KIT+* ILCs did not express detectable *IL17A* (Figure S2B). These results identify αβ CD4+ T cells, not γδ T, MAIT, ILCs or CD8+ T cells, as the primary source of *IL17A* in SpA synovial tissue. In line with previous findings [19], the expression of *IL17F* is not limited to T cells and extend to stroma and endothelial cells (Figure S3). *TNF*, a key driver of SpA pathology, is predominantly expressed by myeloid cells (Figure S4).

**Figure 1.**
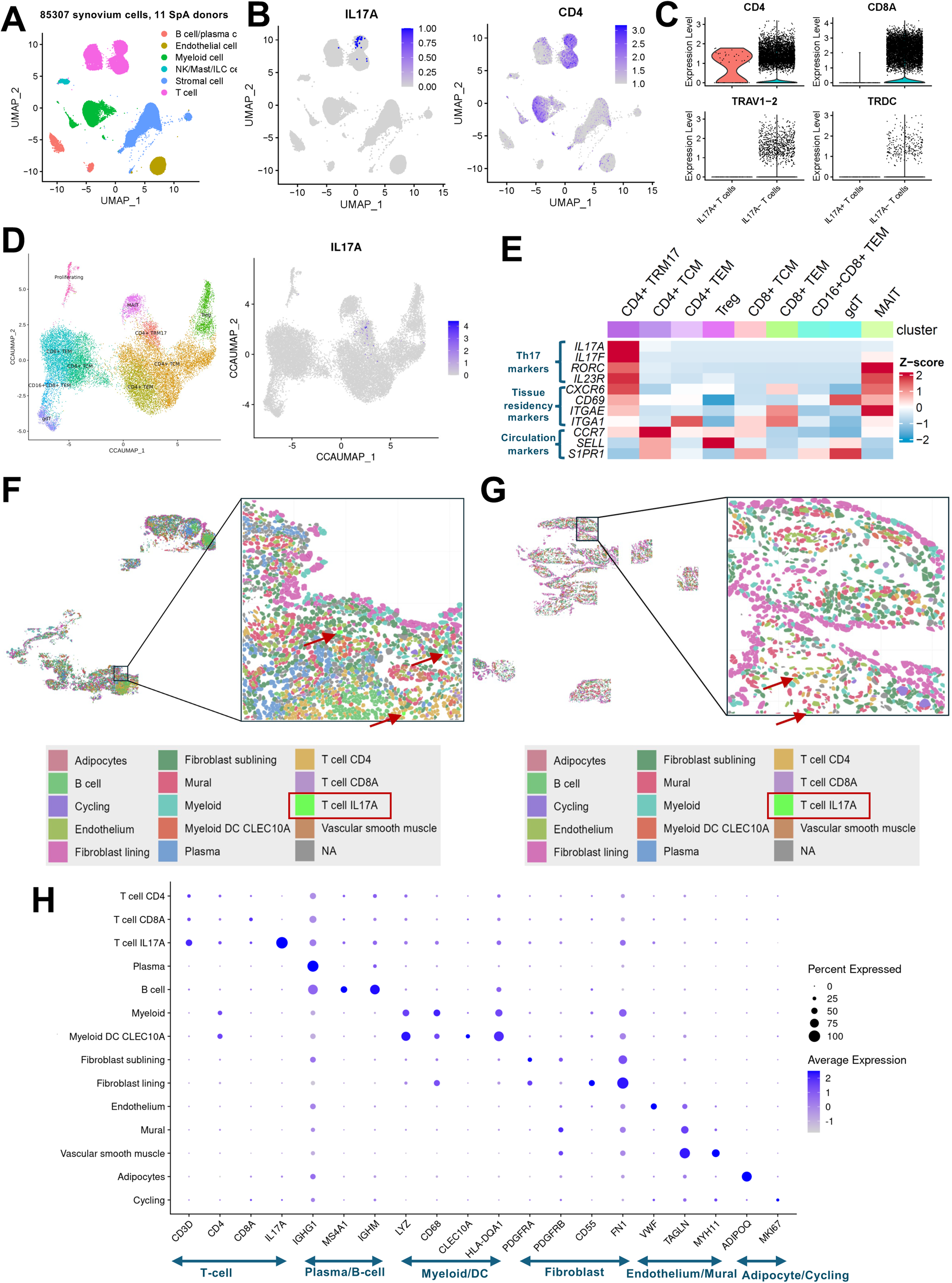
Tissue resident memory CD4+ T cells are the main source of IL17A in SpA joint tissue. (A) UMAP visualization of major cell types of cells from SpA synovial tissue. (B) UMAP visualization of normalized expression of *IL17A* and *CD4*. (C) Violin plot of normalized expression of *CD4, CD8A, TRAV1-2, TRDC* in *IL-17A^+^* and *IL-17A^-^* T cells. (D) UMAP visualization of subsets of T-cells from SpA synovial tissues and normalized expression of *IL17A*. (E) Heatmap of normalized and scaled expression of Th17, tissue residency, and circulation signature markers in different subsets of T cells. CosMx was used for spatial profiling of synovial tissue from four PsA patients. Representative images from two PsA patients are shown in (F) and (G). (H) Expression of cell type marker genes by IL17A+ T cells and other cell populations identified from CosMx data. The “IL17A+ T” subset is excluded from the “CD4+ T” and “CD8+ T” subsets.

To pinpoint the *IL17A* expressing T cell population, we performed sub-clustering analysis for all T cells from SpA synovial tissue. Ten distinct clusters were identified (Figure 1D and Figure S5A). *IL17A* expressing T cells are largely present in the *CD4+* TRM17 population, exhibiting both Th17 (IL17A, IL17F, RORC, IL23R) and TRM (CXCR6, CD69, ITGAE) characteristics (Figure 1E and Figure S5B). Separate analysis of AxSpA and PsA samples confirmed that CD4+ TRM17 cells are the primary source of *IL17A* in both disease states (Figure S6 A-C). Analysis of the AMP rheumatoid arthritis (RA) dataset showed that in RA synovial tissue IL17A was also predominantly expressed by a *CD69+CXCR6+CD4+* TRM17 population (Figure S7 A-C). Notably, CXCR6+CD4+ TRM17 cells have recently been identified to drive the pathology in skin, kidney and brain inflammatory diseases [11,13,14].

To investigate the cellular niche of *IL17A* expressing CD4+ TRM17 cells, we carried out spatial transcriptome analysis on joint synovial tissue from 4 PsA patients using CosMx Spatial Molecular Imaging (Figure 1F and G). We scanned 80 fields of view (FoVs) with a multiplex panel of 1000 genes (CosMx™ Human Universal Cell Characterization RNA Panel). A limitation of this approach was the absence or low expression of many CD4+ TRM cell feature genes identified in our previous scRNA-seq data. However, *IL17A* was successfully detected, enabling the definition of *IL17A*-expressing T-cells which were predominantly localized in the sublining regions (24 of 27 IL17A+ T cells) (Figure 1F and G). We annotated cells based on canonical markers, and identified adipocyte, B, endothelial, sub-lining fibroblast, lining fibroblast, myeloid, CLEC10A+ myeloid, plasma, IL17A+ T, CD4+ T, CD8+ T, and mural cells (Figure 1F-H, Figure S1C and D). Notably, in the analysis shown in Fig. 1H the “IL17A+ T” subset is excluded from the “CD4+ T” and “CD8+ T” subsets.

### Both sublining and lining fibroblasts in SpA joint tissue exhibit an enhanced IL-17 response signature

The induction of a pro-inflammatory state in fibroblasts has been postulated as a critical mechanism underlying IL-17-mediated chronic inflammation. Within the synovium, sub-lining and lining fibroblasts are two distinct subsets that together constitute synovial fibroblasts. We then investigated whether sublining and lining fibroblasts from SpA synovium have been influenced by CD4+ TRM17. To this end, we integrated a publicly available dataset of fibroblasts from healthy synovium[20] with our dataset of SpA synovial fibroblasts, resulting in a combined dataset of 11,173 healthy and 27,975 SpA fibroblasts (Figure 2A). Consistent with previous studies [21,22], we identified THY1+PRG4-sublining and THY1-PRG4+ lining fibroblasts as two major subsets, with MMP1 and MMP3 enriched in the lining fibroblasts (Figure 2A, Figure S9A and B). An IL-17 signalling score (indicative of response to IL-17) was then calculated using a set of genes induced by IL-17 in human synovial fibroblasts[23]. We found both sub-lining and lining fibroblast subsets in SpA exhibited an enhanced IL-17 response signature compared to their healthy counterparts (Figure 2B). Notably, established IL-17-induced effector cytokines and chemokines, such as *CXCL1*, *CXCL8*, *IL6*, and *CCL20*, were consistently upregulated in both sub-lining and lining fibroblast subsets within the SpA synovium. Thus, despite enrichment of CD4+ TRM17 in sublining synovium, both sublining and lining fibroblasts show evidence of IL-17-induced activation.

**Figure 2.**
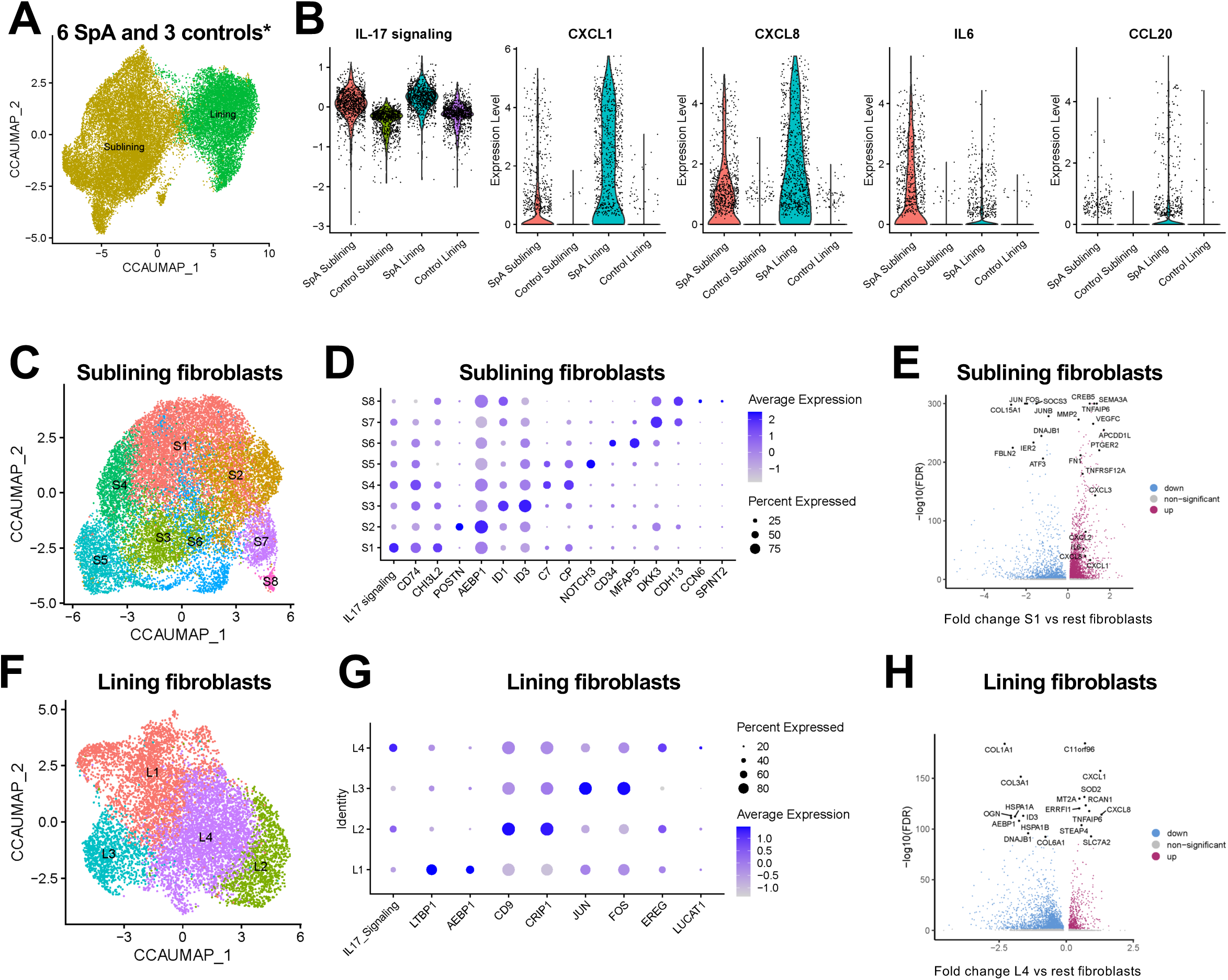
An enhanced IL-17 response signature is present in both sublining and lining fibroblasts from SpA synovium. (A) UMAP visualization of fibroblast subsets of combined cells from SpA and healthy synovial tissues. (B) Violin plot of IL-17 signalling score and normalized expression of four selected IL-17 signalling genes in different fibroblast subsets in SpA and healthy synovium. The data for control synovium (n=3) were sourced from GSE216651. (C) UMAP visualization of subsets of sub-lining fibroblast cells from SpA synovium. (D) Relative expression of marker genes of different sub-lining fibroblast subsets from SpA synovium. (E) Volcano plot of differentially expressed genes in S1 sub-lining fibroblast cells compared to other sub-lining fibroblast cells in SpA synovium (FDR < 0.05). (F) UMAP visualization of subsets of lining fibroblast cells from SpA synovium. (G) Relative expression of marker genes of different lining fibroblast subsets from SpA synovium. (H) Volcano plot of differentially expressed gene in L4 lining fibroblast cells compared to other lining fibroblast cells in SpA synovium (FDR < 0.05).

We then asked if the enhanced IL-17 signature in SpA synovium fibroblasts is contributed by certain subset(s) within the sub-lining and lining fibroblast populations. To this end, we conducted a detailed analysis of cell states within these two populations separately (Figure 2C and F). In the sub-lining fibroblasts, among eight identified subsets, a cluster (S1) marked by the expression of *CD74* and *CHI3L2* exhibited the highest IL-17 signalling score (Figure 2D). This subset was also characterized by the upregulation of pro-angiogenic genes, including *SEMA3A* and *VEGFC* (Figure 2E). In the lining fibroblast compartment, the L4 cluster displayed the strongest IL-17 signalling signature (Figure 2G) and was enriched in the expression of genes associated with oxidative stress and inflammatory responses, such as *SOD2*, *MT2A*, *RCAN1*, and *TNFAIP6* (Figure 2H).

### CLEC10A+ dendritic cells are potential antigen presenting cells for CD4+ TRM17 cells

Tissue-resident memory T cells typically require antigen presenting cells (APCs) for their generation and effector function. It has been well established that myeloid cells especially dendritic cells (DCs) are the key APCs to support T cell function. To identify the myeloid cellular subset that serves as APC for CD4+ TRM17 cells, we extracted and re-clustered the myeloid cells from SpA tissue (Figure 1A) and identified ten transcriptionally distinct myeloid subsets: plasmacytoid dendritic cells (pDC), *CLEC9A*+ DC, *CLEC10A*+ DC, *TNF*+ macrophage cluster 1 (M1), *TNIP3*+ macrophage cluster 2 (M2), *EGR2+* macrophage cluster 3 (M3), *GIMAP*+ macrophage cluster 4 (M4), *C9*+ macrophage cluster 5 (M5), *PRODH2*+ macrophage cluster 6 (M6), and *LYVE1*+ macrophage cluster 7 (M7) (Figure 3A). We then searched for potential APCs for CD4+ TRM17 using cell-cell communication analysis for all ten myeloid subsets and *IL17A*+ T cells (Figure 3B). The strongest interaction with *IL17A*+ cells was observed with *CLEC10A*+ DCs, which expressed high levels of ligands for CD28 (*CD80* and *CD86*), produced a chemokine (*CCL20*) known to be chemotactic for Th17 cells through CCR6 and were enriched for the key cytokine for the generation of tissue resident memory T cells (*TGFB1*, encoding TGF-β).

**Figure 3.**
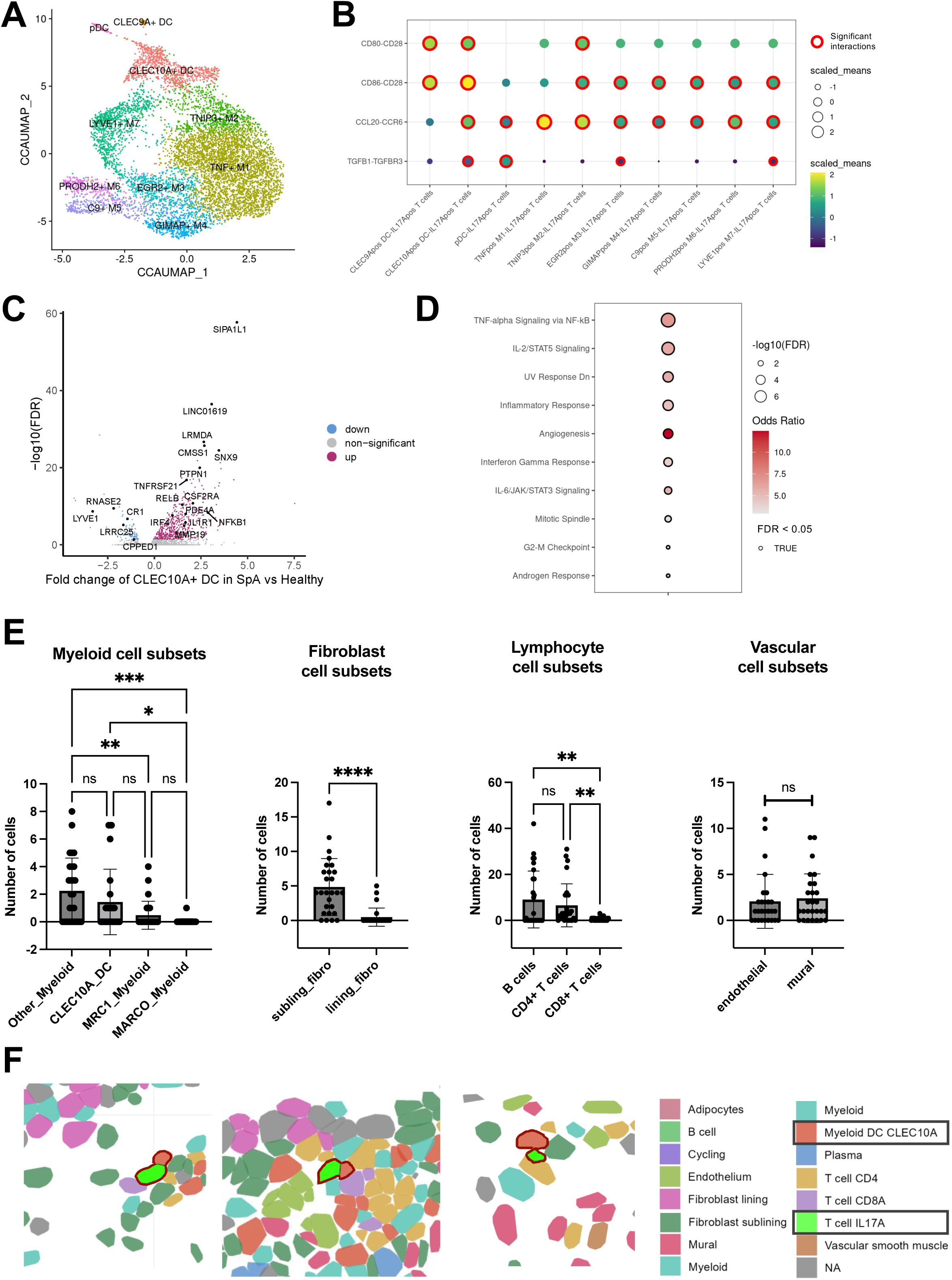
Cell-cell communication and single-cell spatial analysis support the interaction between CLEC10A+ dendritic cells and TRM17. (A) UMAP visualization of subsets of myeloid cells from SpA synovial tissues. (B) Cell-cell interactions between different myeloid subsets and IL17A+ T cells in SpA synovium inferred by CellPhoneDB. (C) Volcano plot of differentially expressed gene in CLEC10A+ DC cells in SpA synovium compared to healthy synovium (FDR < 0.05). (D) Enriched pathway in CLEC10A+ DC cells (All up-regulated genes were used for enrichment analysis using MSigDB database). (E) The number of myeloid, fibroblast, lymphocyte, and vascular cell subsets located within 50µm of *IL17A+* T cells was quantified. (F) Three representative CosMx images of SpA synovium showing interaction between CLEC10A+ DC cells and IL17A+ cells (highlighted in red outline). Statistical significance was assessed using one-way ANOVA (for myeloid and lymphocyte subsets) or the paired two-tailed Student’s t-test (for fibroblast and vascular subsets) (* P ≤ 0.05, ** P ≤ 0.01, *** P ≤ 0.001, **** P ≤ 0.0001). Median +/-SEM is shown in (E).

Activated T cells have been shown to act on myeloid cells and induce their activation [24] [25]. Thus, if *CLEC10A*+ DC cells have served as the APC for CD4+ TRM17 cells, they should have received reciprocal signals from these cells and show signs of activation. To look for evidence of *CLEC10A*+ DC activation in SpA synovium, we carried out a differential gene expression analysis which revealed 541 upregulated and 90 downregulated genes in *CLEC10A*+ DCs in SpA versus healthy synovium (Figure 3C). Notably, genes associated with an inflammatory response (*RELB*, *IRF4*, and *NFKB1*) were significantly increased. Pathway enrichment analysis of these upregulated genes using the MSigDB database revealed significant enrichment in several pro-inflammatory pathways, including TNFα signalling, IL-2/STAT5 signalling, IFNγ response, and IL-6/JAK/STAT3 signalling (Figure 3D). However, pro-inflammatory pathways were also elevated in TNF+ M1 and LYVE1+ M7 cells from SpA patients (Figure S10). Therefore, we cannot determine whether the enrichment of pro-inflammatory pathways in SpA *CLEC10A*+ DCs is due to T cell interactions or the joint’s inflammatory environment. Based on the hypothesis that antigen-presenting cells (APCs) for IL17A+ T cells would have encountered IL-17, we analyzed the expression of genes known to be induced by IL-17A in monocytes or macrophages (MMP9, CCL4, CCL5, CSF2, IL3, IL9) [26,27]. We observed increased expression of MMP9 and CSF2 in CLEC10A+ dendritic cells (DCs), but a greater number of genes were enriched in TNF+ M1 and TNIP3+ M2 macrophages (Figure S11). Lastly, we sought evidence of direct physical cell interactions between CLEC10A+ dendritic cells (DCs) and IL17A+ T cells in our spatial transcriptomic datasets. Quantification of cell proximity revealed that CLEC10A+ DCs were one of the top two subsets significantly enriched in close proximity to IL17A+ T cells (Figure 3E). Representative examples of CLEC10A+ DC and IL17A+ T cell interactions are shown in Figure 3F. We also observed that IL17A+ T cells are more frequently in proximity to sublining fibroblasts than lining fibroblasts, and to B cells and CD4+ cells than CD8+ cells. Taken together, these data suggest that *CLEC10A*+ dendritic cells are activated in SpA joints and are the potential APCs for CD4+ TRM17.

### T-cell receptor engagement and cytokine stimulations co-drive the production of IL-17A/F by in vitro generated CD4+ TRM17 - like cells

We then asked what signals induce the production of IL-17A by CD4+ TRM17 cells. Due to the difficulties in obtaining adequate number of CD4+ TRM17 from synovial tissue for functional studies, we developed an *in vitro* model to induce CD4+ TRM17-like cells. It has been shown that following immunization effector Th17 cells can give rise to lung CD4+ TRM17 cells which are maintained in the tissue by the survival cytokine IL-7 following the clearance of antigens[12]. Accordingly, we generated a two-phase *in vitro* model to generate CD4+ TRM17-like cells (Figure 4A). In phase I, circulating blood memory CD4+ T cells from patients with SpA were stimulated with anti-CD3 antibody, anti-CD28 antibody and Th17 expansion cytokines (IL-1β and IL-23) to generate effector Th17 cells. These cells were then cultured with TGF-β (to model the tissue niche) and IL-7 (to provide the survival signal) in the absence of T cell receptor stimulation in phase II. Using this model, we were able to induce IL-17A+CXCR6+CD4+ TRM17-like cells which were largely absent in the blood memory compartment (Figure 4B and C). We then assessed the expression of TRM markers using our assay in comparison to a 6-day Th17 expansion assay (anti-CD3/CD28, IL-1β, IL-23). While the CD4+ TRM17 assay slightly improved IL-17A+ cell induction, it significantly increased the percentage of Th17 cells expressing CXCR6, CD49a, CD103, and CD69 (Figure S12A, B). This was confirmed at the RNA level by qPCR (Figure S12C). Taken together, these data suggest that our assay successfully induces Th17 cells with TRM-like features in vitro.

**Figure 4.**
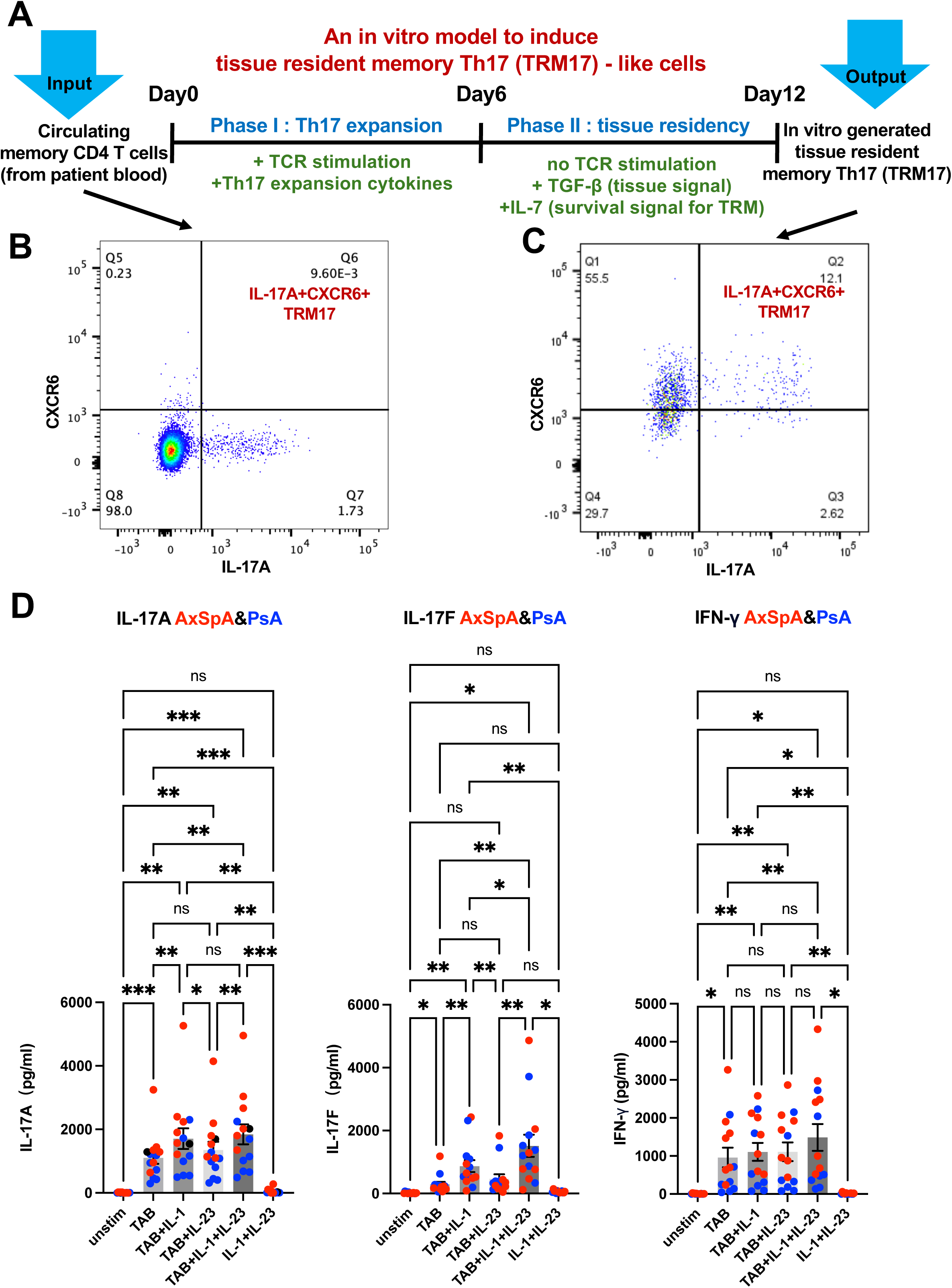
T-cell receptor engagement and cytokine stimulations co-drive the production of IL-17A/F by in vitro generated TRM17 - like cells. (A) Schematic representation of the in vitro TRM17 model. Assessment of IL-17A+CXCR6+ TRM17 in circulating memory CD4+ T cells (B) and induced TMR17 cells (C). (D) The secretion of IL-17A, IFN-γ and IL-17F by TRM-like cells stimulated with vehicle control, cytokines (IL-1β and IL-23), T-cell activation beads (TAB) or the combinations of cytokine and TAB. Data from 7 AxSpA (red) and 7 PsA (blue) patients are shown. Statistical significance was assessed using one-way ANOVA (* P ≤ 0.05, ** P ≤ 0.01, *** P ≤ 0.001). Median +/-SEM is shown in (D).

Whilst IL-23 blockade has proved highly efficacious for skin psoriasis, the effects on AxSpA and psoriatic arthritis have been less impressive [28–31]. This led us to investigate the role of IL-23 in IL-17A production by CD4+ TRM17-like cells. Figure 4D shows that T cell receptor engagement using anti-CD3 and anti-CD28 T cell activation beads (TAB), rather than cytokine stimulation (IL-1β and IL-23), induces IL-17A production in restimulated CD4+ TRM17-like cells from SpA patients. Furthermore, the addition of IL-23 to either TAB or TAB + IL-1β conditions did not significantly enhance IL-17A production. Interestingly, in the presence of TAB and IL-1β, IL-23 significantly increased the secretion of IL-17F. This suggests that the lack of effect of IL-23 on IL-17A production is not due to the pre-existence of IL-23 in the culture media. Consistent with this, IL-23 blockade did not reduce IL-17A secretion in either the TAB alone or TAB + IL-1β conditions (Figure S13). IL-1β blockade has been tested in AxSpA and PsA with mixed results. We observed an IL-17-specific enhancement by IL-1β, but not IFN-γ, either in TAB or TAB + IL-23 conditions. A similar pattern was observed when AxSpA and PsA samples were analyzed separately (Figure S14), as well as in CD4+ TRM17-like cells generated from healthy controls (Figure S15). We also performed a similar experiment using cells generated from a 6-day Th17 expansion assay (anti-CD3/CD28, IL-1β, IL-23) (Figure S16). Like the CD4+ TRM17 assay, IL-17A secretion was predominantly driven by TAB and IL-1β. Surprisingly, IL-17F was only induced when both TAB and cytokines (IL-1β or IL-23) were present. Taken together, these data suggest that both T cell receptor engagement and cytokine stimulation co-drive the production of IL-17A/F by in vitro generated CD4+ TRM17-like cells.

### Identification of BRD1 as a novel epigenetic regulator of Th17/CD4+ TRM17 cells

The unique epigenetic profile of CXCR6+Th17 cells reported in the murine multiple sclerosis model [14] drove us to hypothesize that epigenetic regulators could contribute to the generation of CD4+ TRM17 cells. To test this hypothesis, we screened a library of 38 epigenetic inhibitors using cells from SpA patients. Due to the low-throughput nature of the CD4+ TRM17 model, we performed the initial screen using a Th17 expansion assay and discovered three inhibitors (Bromosporine, PFI-1 and OF-1) that suppress Th17 responses in both intracellular cytokine staining (ICS) and IL-17 enzyme-linked immunosorbent assays (ELISA) (Figure 5A). We deprioritized the first two inhibitors as Bromosporine is a pan-bromodomain inhibitor with multiple targets and the target of PFI-1, BET Bromodomain, has already been shown to regulate Th17 cells[32,33]. In contrast OF-1, an inhibitor for the bromodomain and PHD finger-containing protein (BRPF) family [34], has not been implicated previously in Th17 response and thus was chosen for further investigation. To this end, we first used siRNA to screen all three members in the BRPF family and found BRD1 silencing inhibited IL-17A production (Figure 5B). A similar trend was observed in BRPF1-silenced cells. Silencing efficiency was greater than 50% for all three genes (Figure S17A), a level generally considered acceptable for primary immune cell experiments. Further transcriptome analysis showed that knockdown of BRD1, but not BRPF1, reduced both Th17 marker genes (*IL17A, IL17F, RORC and IL23R*) and TRM markers (*CXCR6 and ITGA1*) (Figure 5C). Taken together these data suggest a potential role of BRD1 in CD4+ TRM17 cell generation.

**Figure 5.**
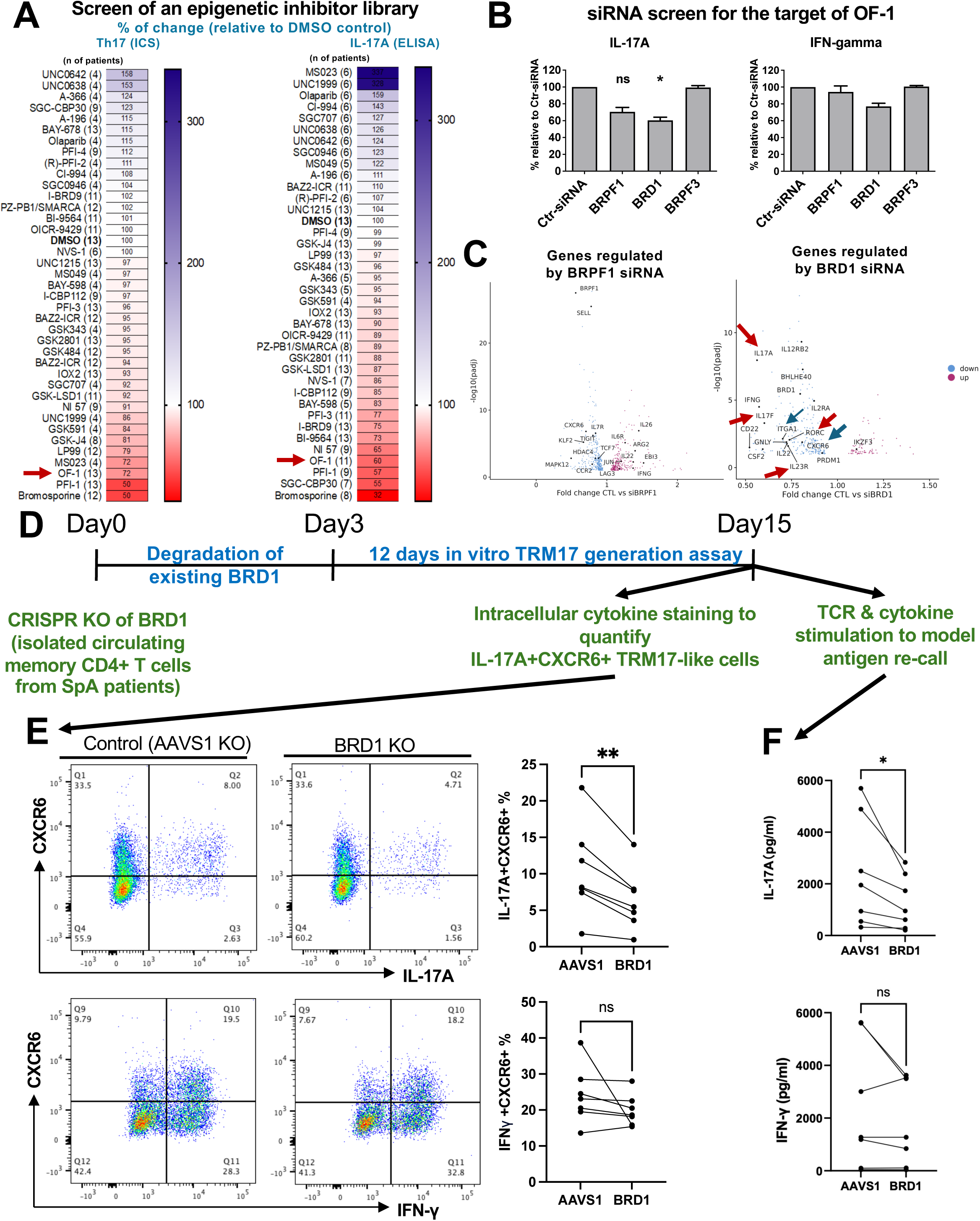
The epigenetic regulator BRD1 plays a role in TRM17 generation. A library of epigenetic inhibitors was screened using CD4+ T cells from SpA patients. Depending on the amount of blood obtained, a range of 4-13 donors were recruited for different inhibitors. The changes of Th17 frequency (based on IL-17A intracellular cytokine staining) and the level of IL-17A secretion (ELISA) relative to vehicle control (DMSO) are shown (A). (B) The respective influence of siRNAs for BRPF1, BRD1 and BRPF3 on IL-17A secretion by CD4+ T cells stimulated with T cell activation beads (TAB) and Th17-promoting cytokines (IL-1β and IL-23) (data from 3 SpA patients). (C) The transcriptome changes induced by the siRNA silencing of BRPF1 and BRD1 (data from 6 SpA patients). (D) Schematic representation of BRD1 knockout for the in vitro TRM17 model. (E and F) Effect of BRD1 knockout on the generation of TRM17-like cells and their IL-17A production upon antigen re-call (data from 7 SpA patients). The P-value was assessed using the Friedman test for (B) and paired student’s T test for (E and F) (* P≤ 0.05; ** P≤ 0.01). Median +/- is shown in (B).

We then used our CD4+ TRM17 in vitro model to investigate the role of BRD1 in the generation of TRM17-like cells. As the transient nature of siRNA-mediated gene suppression is not suitable for the 12 days CD4+ TRM17 model, we deployed clustered regularly interspaced short palindromic repeats (CRISPR) technology to stably knockout BRD1 (Figure 5D) and achieved over 50% knockout efficiency (Figure S17B). The BRD1-KO CD4+ T cells were rested for three days (allowing the degradation of endogenous BRD1 protein) before being used as the input for CD4+ TRM17 generation model. Figure 5E shows that, in comparison to AAVS control guide RNA, BRD1 knockout significantly inhibited the generation of CXCR6+IL-17A+ TRM17-like cells (Figure 5E). No changes in cell viability or proliferation were observed. Additionally, the restimulation of TMR17-like cells induced a lower level of IL-17A secretion in BRD1-knockout cells than the AAVS control cells (Figure 5F). Similar results were observed for IL-17F (Figure S17). In contrast to IL-17A and IL17-F, the percentage of IFN-γ+CXCR6+ cells and the secretion IFN-γ were not affected by BRD1 KO (Figure 5E and F).

## Discussion

In this study, we show that CXCR6+ tissue resident memory CD4+ T cells are the main source of IL-17A in SpA joint tissue. Despite the predominant localization of CD4+ TRM17 cells in the sublining regions, both sublining and lining fibroblasts in SpA joints exhibit an enhanced IL-17 response signature suggesting the diffusion of IL-17A within the tissue. We also show that T-cell receptor engagement and cytokine stimulations co-drive the production of IL-17A/F by in vitro generated TRM17 - like cells. Lastly, we identify BRD1 as a novel Th17 regulator that contributes to the generation of CD4+ TRM17-like cells in vitro.

Previous studies in SpA synovial fluid (SF) identified γδ T cells, MAIT cells, iNKT and CD8+ tissue-resident memory T cells as key sources of IL-17 [3–10]. Our findings indicate CD4+ TRM17 cells as the primary source of IL-17 in SpA synovial tissue (ST). This discrepancy suggests that distinct inflammatory drivers are present in SF versus ST. The relationship between SF and ST inflammation within IL-17 pathology and beyond warrants further investigation. Our SpA synovial tissue dataset makes a significant contribution to existing knowledge. In addition to demonstrating CD4 TRM IL-17A and IL-17F production, we also highlight the activation and function of various non-T cell populations within the SpA synovium. Specifically, we observe TNF production primarily by myeloid cells (mainly macrophages). Furthermore, we detect the activation of fibroblasts and *CLEC10A+* DCs in these AxSpA patients, compared to healthy controls, using infrapatellar fat pad and synovium tissues studied elsewhere.

Importantly, we found the *in vitro* generated CD4+ TRM17-like cells produced IL-17A in response to T cell receptor (TCR) engagement rather than cytokine stimulation. In the presence of IL-1β, IL-23 enhanced the TCR stimulation induced production of IL-17F but not IL-17A, suggesting that cytokine stimulation has a greater influence on IL-17F production. In line with this, IL-17F has been shown to be the dominant IL-17 isoform produced by cytokine-activated innate lymphocytes (MAIT cells, γδ T cells and ILC3s) [35,36]indicating a closer association of IL-17F with the innate response in lymphocytes. We also observed that IL-1β enhanced IL-17A/F production in the presence of TCR engagement, and this enhancement occurred both with and without IL-23.

We acknowledge the inherent limitations of this study. Our sample size is relatively small, largely due to the costs of scRNA sequencing and the availability of tissue. More than half of the samples came from patients with long disease duration (>10 years). While we cannot be certain that our findings fully represent early-stage SpA, we did detect CD4 TRM17 IL-17 production in samples from patients with a shorter disease duration (<5 years). A larger cohort, stratified by disease duration (and therapy), would be necessary to further confirm and expand upon our findings. In addition, we used human infrapatellar fat pad and synovial tissues as controls for the study of fibroblasts and myeloid cells. A comparison with synovium from healthy knee, hip, or spine should be conducted in the future to confirm our findings.

In summary, we demonstrate for the first time that the CD4+ TRM17 population is the main source of IL-17A in SpA synovial tissue. In addition, our data suggest that targeting CD4+ TRM17, the “factory” of IL-17A, has the potential to induce long-term remission, encouraging future efforts to develop new therapies to deplete CD4+ TRM17 cells in SpA.

## Acknowledgements

We thank the patients for agreeing to participate in research and the Birmingham Tissue Analytics facility for generating the CosMx data. This work was funded by a Versus Arthritis career development award to LC 22053. FL, PB, CB, HU were supported by the National Institute for Health Research (NIHR) Oxford Biomedical Research Centre (BRC). The views expressed are those of the author(s) and not necessarily those of the NHS, the NIHR or the Department of Health.

## Competing interests

This work has been partially funded by J&J to the Cartography Consortium. LC has received research support from Novartis. PB has received research support from Regeneron, Benevolent AI, Novartis and GSK. H.H.U. has received research support or consultancy fees from J&J, UCB Pharma, Eli Lilly, BMS Celgene, GSK.

## Research Ethics Approval

Venous blood was obtained under protocols approved by the Oxford Research Ethics committee (ethics reference number 06/Q1606/139). Synovial tissue were obtained under protocols approved by the Oxford Research Ethics committee (ethics reference number 06/Q1606/139), South Birmingham Research Ethics committee (ethics reference number 14/WM/1109) and Ethics Committee of Shanghai Sixth People’s Hospital (2024-KY-132).

## Data availability

Data has been deposited at GEO (GSE290921) and will be released following the publication of the work.

## Notes

### Summary of Updates

Changes have been made in all figures.

