## Supplementary material for "CD4+ tissue-resident memory Th17 cells are a major source of IL-17A in Spondyloarthritis synovial tissue": N/A

**Supplementary Table 1: Demographics of patients studied**

| | AxSpA patients recruited for scRNA-seq* | PsA patients recruited for scRNA-seq | PsA patients recruited for spatial transcriptomics | AxSpA patients recruited for cellular assays | PsA patients recruited for cellular assays |
|---|---|---|---|---|---|
| Number | 5 | 6 | 4 | 10 | 8 |
| Female/Male | 1/4 | 3/3 | 1/3 | 4/6 | 4/4 |
| Age (mean in years) | 61.8 | 46.1 | 51 | 50.8 | 48.1 |
| Disease duration (mean in years) | 23.2 | 9.3 | NA | 18.9 | 9.7 |
| Current biological therapy | 1** | 1*** | 4**** | 3 | 2 |
| Current synthetic DMARD therapy | 0 | 3 | 0 | 0 | 8 |
| Current steroid therapy | 0 | 0 | 0 | 0 | 0 |
| HLA-B27 positive/negative | 4/1 | NA | NA | 7/3 | NA |
| CRP mean (mg/L) | 13.3 | 13.0 | 15.8 | 6.7 | 2.1 |
| BASDAI mean | 3.4 | NA | N/A | 6.2 | NA |
| Swollen joint count mean (/66) | NA | 3.8 | 8.5 | NA | 2.5 |
| Tender joint counts mean (/66) | NA | 10 | 14.2 | NA | 11.6 |

*all had radiographic AxSpA, 4 samples from hip, one from knee
** 1 current TNFi (etanercept), 4 previous TNFi therapy
*** 1 current TNFi, 3 previous TNFi therapy
**** 4 current TNFi (adalimumab)

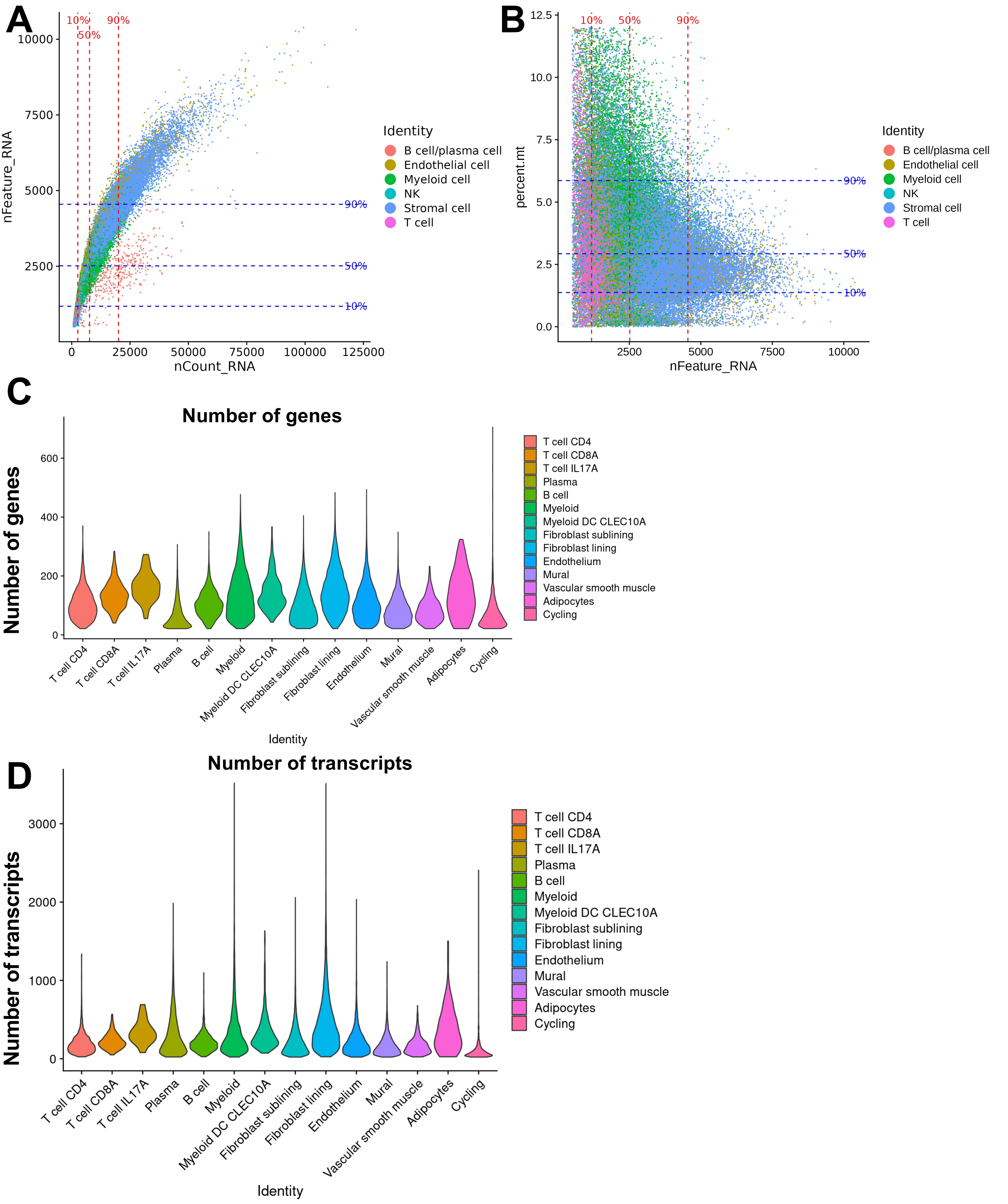


**Figure S1. Quality control (QC) plots for SpA scRNA-seq and spatial transcriptome dataset.** (A) Distribution of the number of detected features (nFeature_RNA) versus total RNA count (nCount_RNA).(B) Distribution of the percentage of mitochondrial genes (percent.mt) versus the number of detected features (nFeature_RNA). Cells are color-coded by cell identity: B cell/plasma cell, endothelial cell, myeloid cell, NK cell, stromal cell, and T cell. The red dashed vertical lines in panel A represent the 10%, 50%, and 90% quantiles of the nCount_RNA, while the blue dashed horizontal lines indicate the 10%, 50%, and 90% quantiles of the nFeature_RNA. In panel B, the 10%, 50%, and 90% quantiles for percent.mt are indicated by the horizontal blue dashed lines. For spatial RNA profiling datasets, number of genes and transcripts per cell subsets is shown in (C) and (D) respectively.

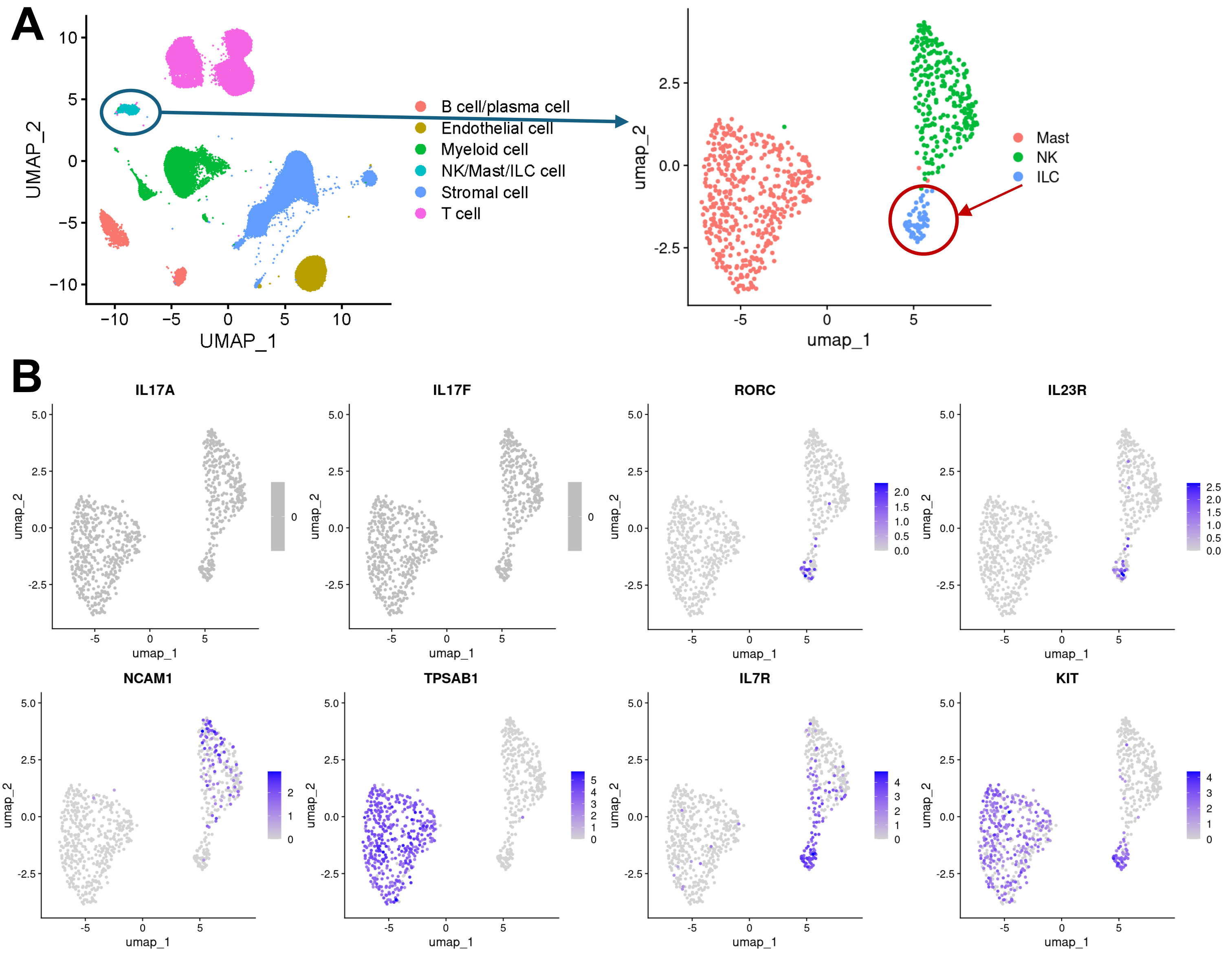


**Figure S2. SpA joint tissue ILCs, NK and Mast cells do not produce detectable *IL17A or F*.** (A) UMAP visualization of re-clustered cells from NK/Mast/ILC population from cells shown in Figure 1A. (B) UMAP visualization of normalized expression of selected marker genes.

A

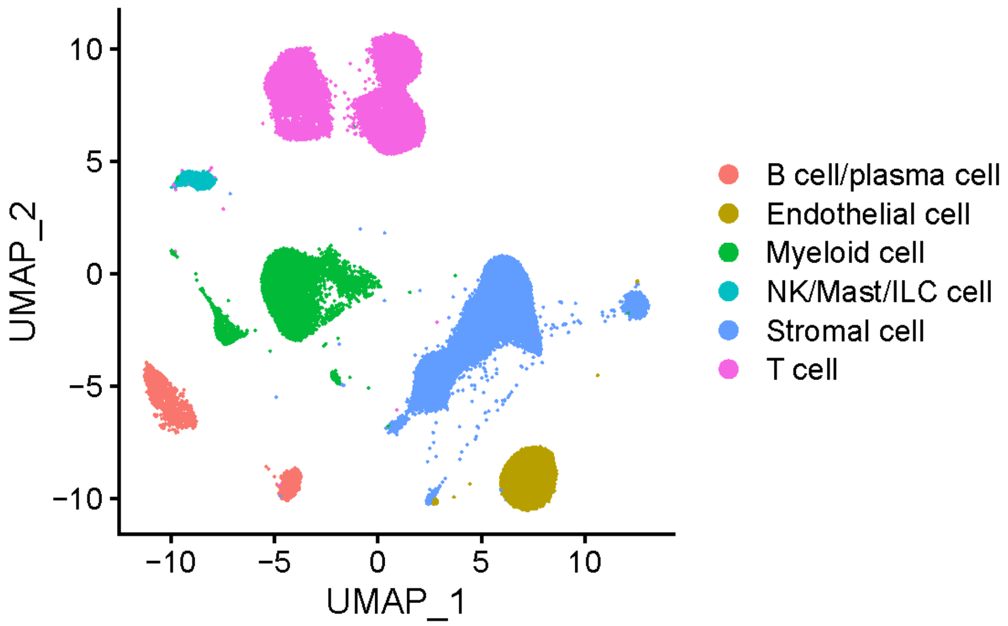


B

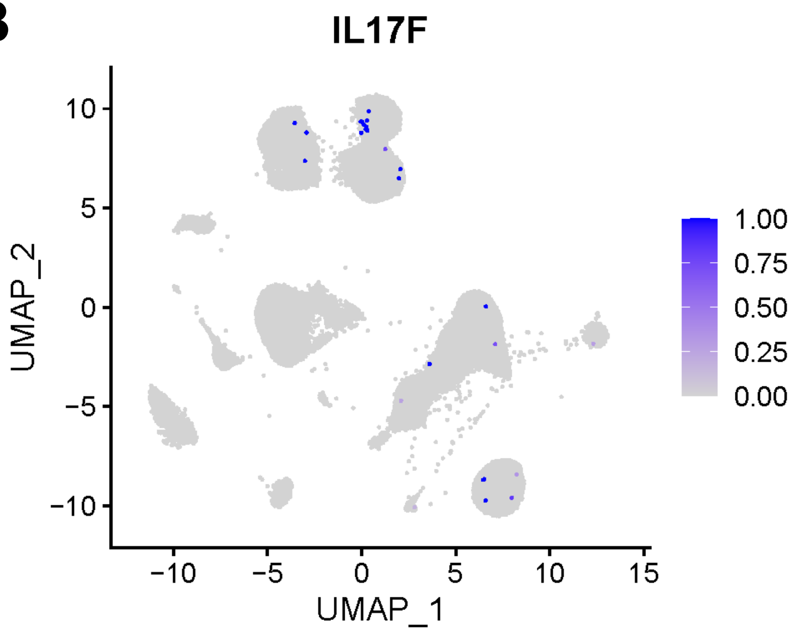


**Figure S3. Multiple SpA joint tissue cell types express *IL17F*.** UMAP visualization of normalized expression of *IL17F* by SpA synovial tissue cells.

**A**

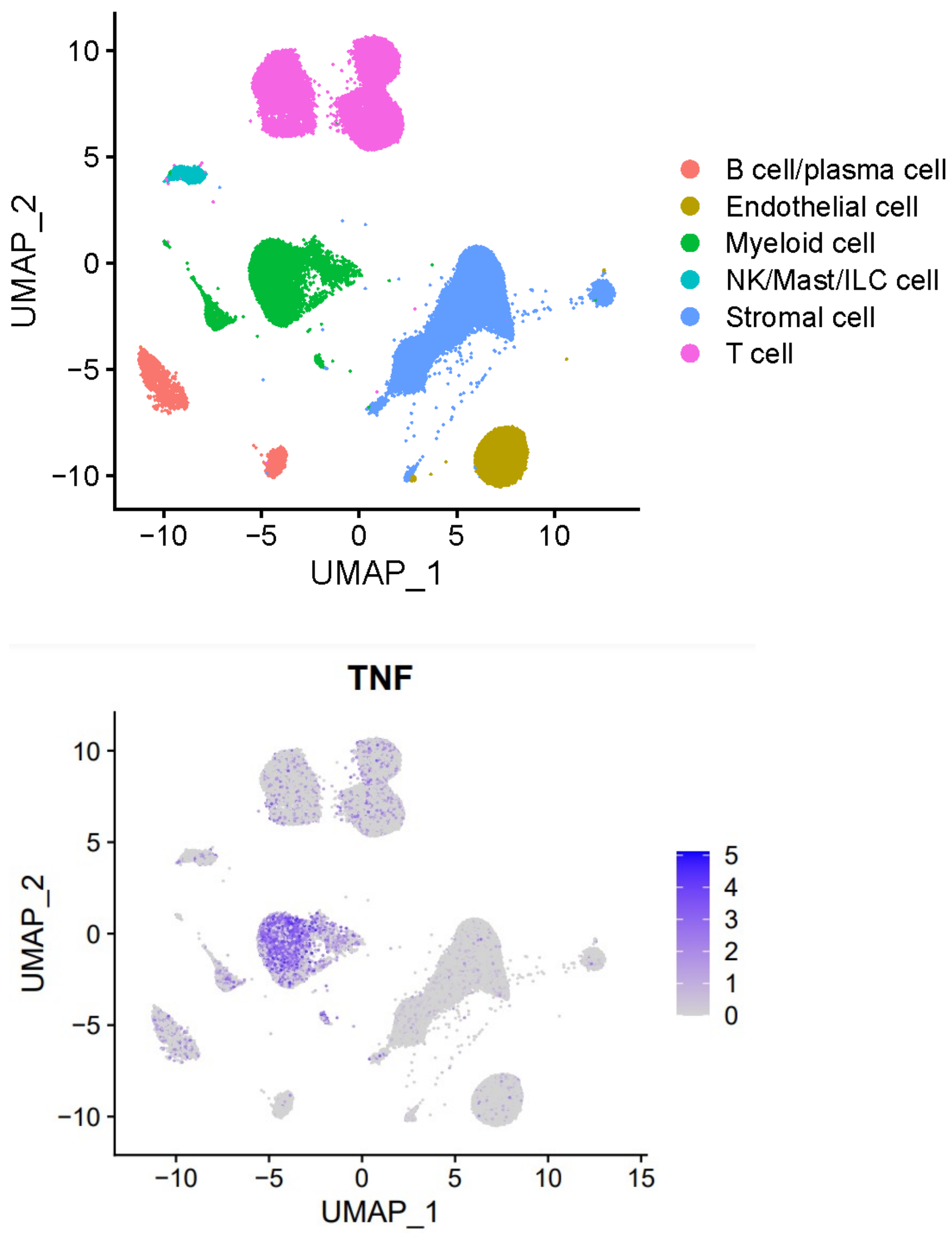


**B**



**Figure S4. Within SpA synovial tissue *TNF* is predominantly expressed by myeloid cells.** UMAP visualization of normalized expression of *TNF* by SpA synovial tissue cells.

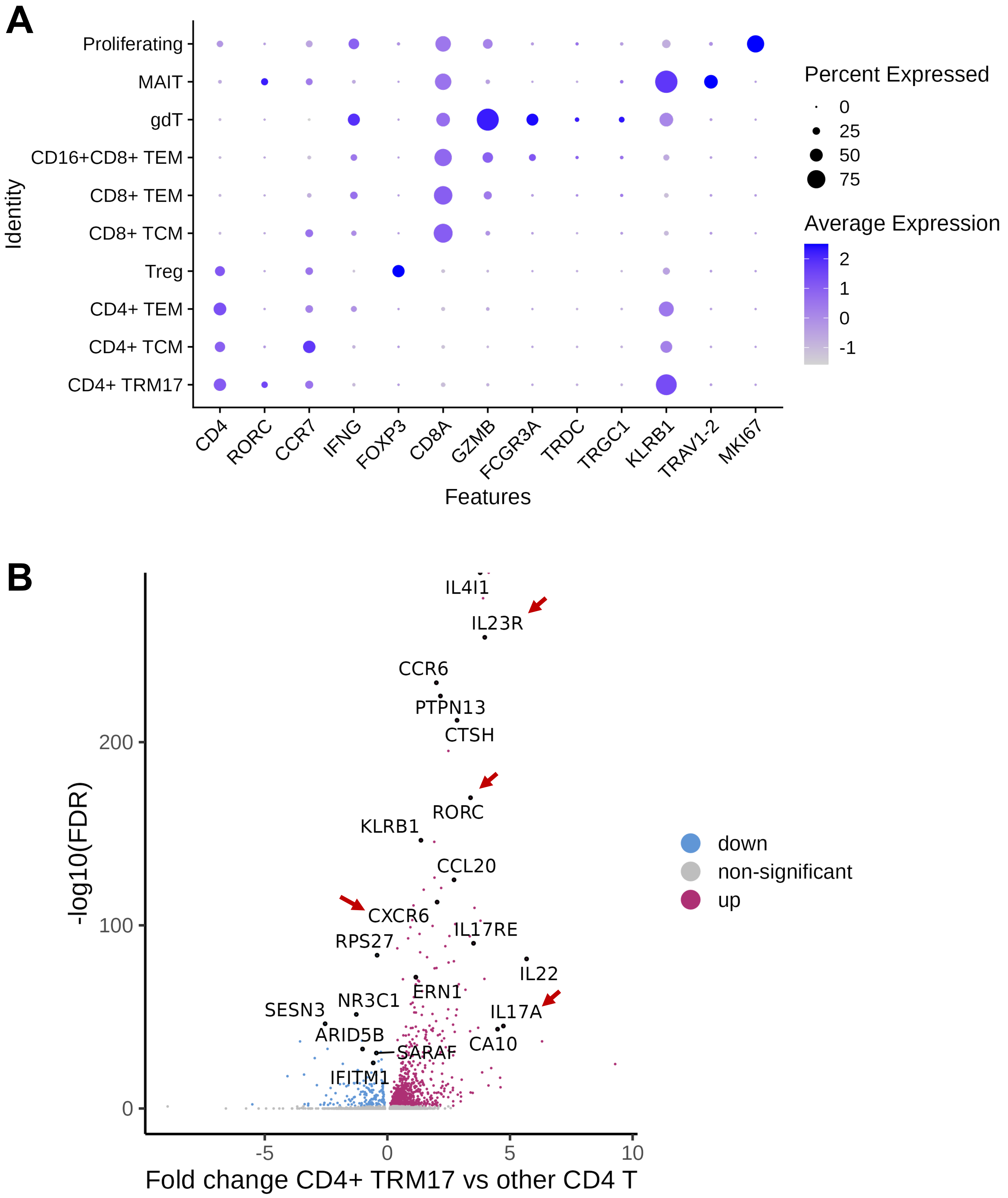


**Figure S5. Sub-clustering of SpA T-cells identifies a *CD4*+ TRM17 population in SpA synovial tissue.** (A) Relative expression of maker genes of different T-cell subsets from SpA synovium. (B) Volcano plot of differentially expressed gene in CD4+ TRM17 cells compared to other non-Treg CD4 T cells (FDR < 0.05).

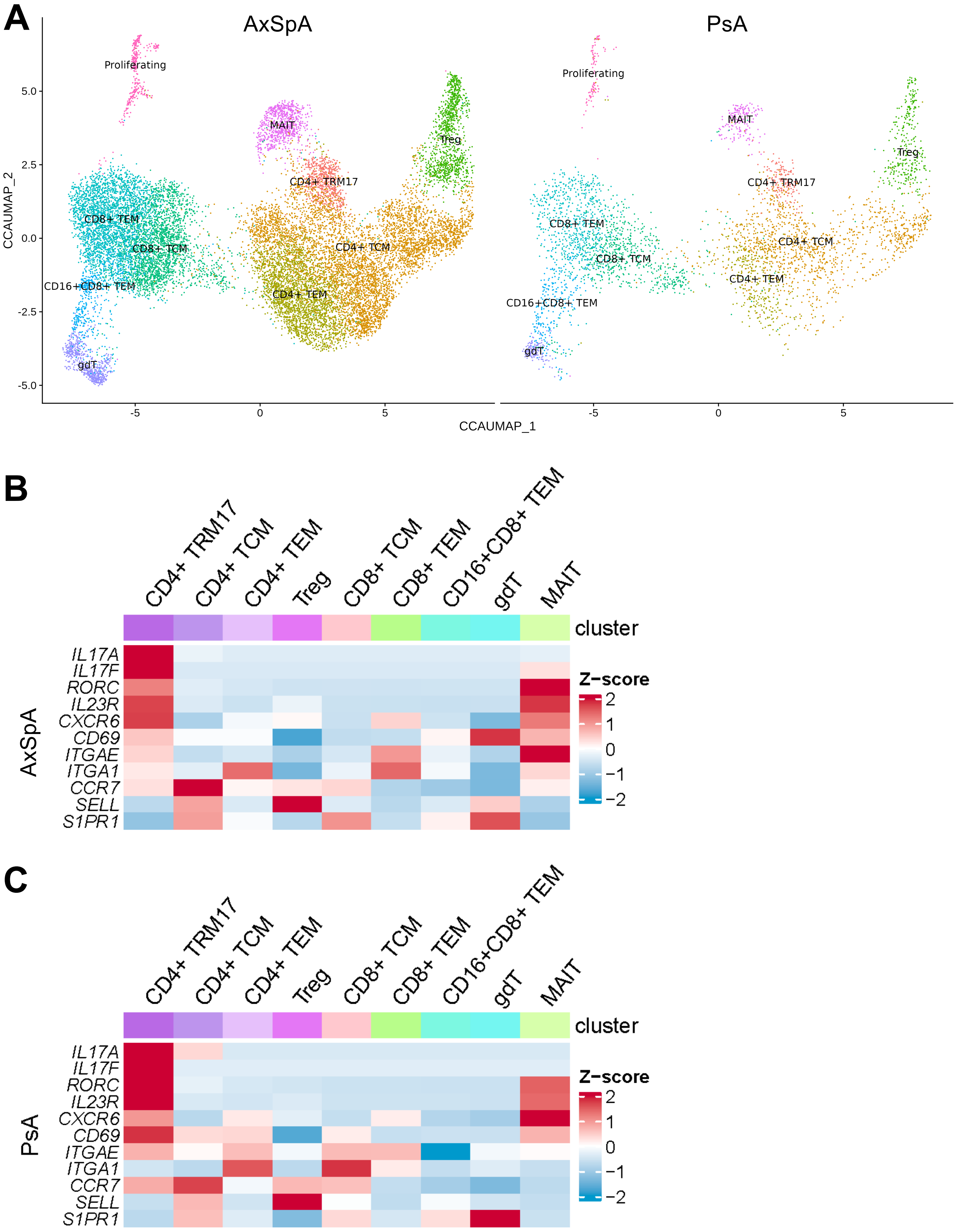


**Figure S6. *IL17A* is predominantly expressed by CD4+ TRM17 population in both AxSpA and PsA synovial tissue.** (A) UMAP visualization of subsets of T-cells from synovial tissues from AxSpA and PsA patients. Heatmap of normalized and scaled expression of Th17, tissue residency, and circulation signature markers in different subsets of T cells from AxSpA (B) and PsA (C).

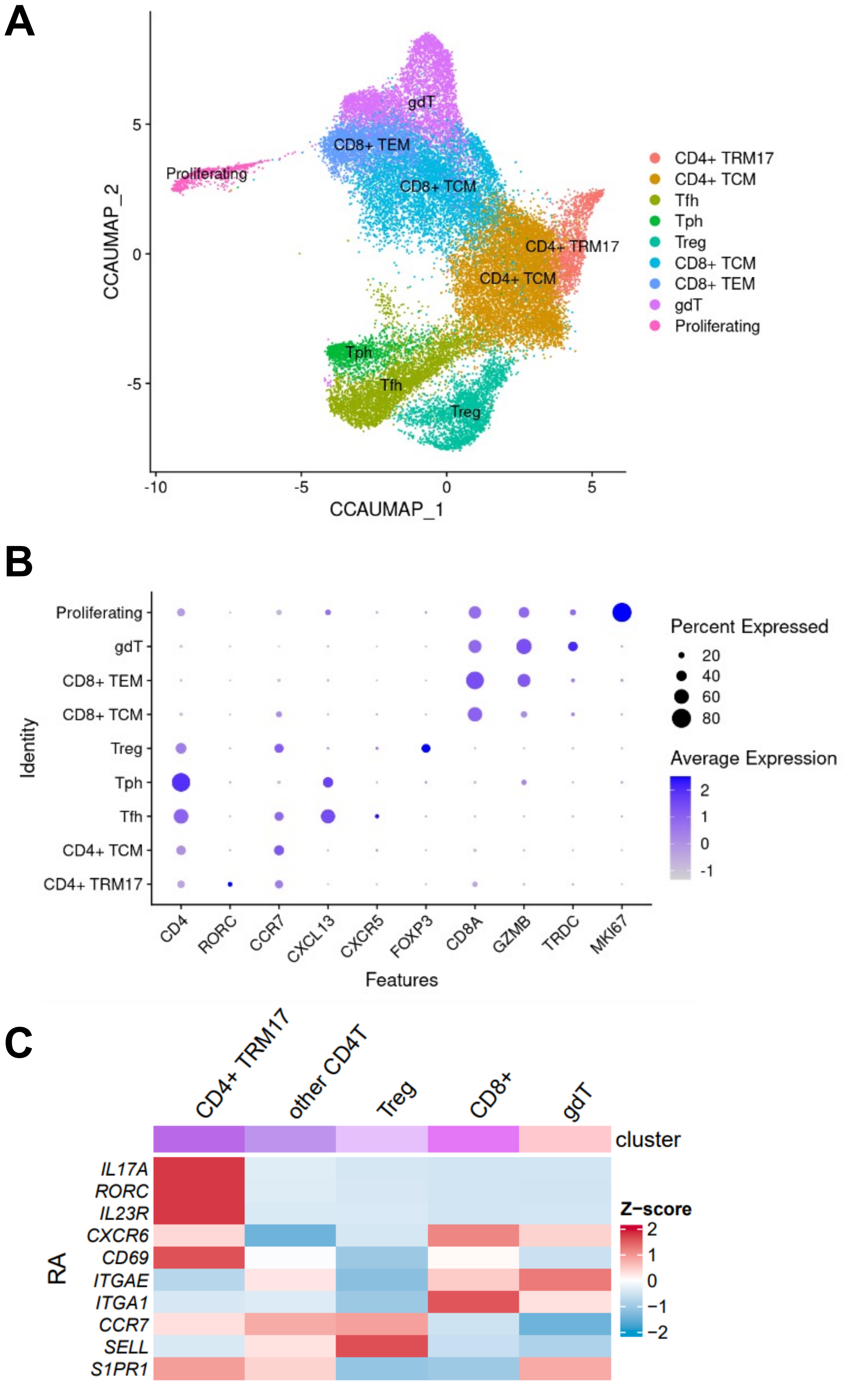


**Figure S7. *IL17A* is predominantly expressed by CD4+ TRM17 population in RA synovial tissue.** T-cell scRNA-seq data was sourced from the AMP RA Phase II study (Zhang F et al. Nature 2023). (A) UMAP visualization of subsets of T-cells from SpA synovial tissues. (B) Relative expression of maker genes of different T-cell subsets from RA synovium. (C) Heatmap of normalized and scaled expression of Th17, tissue residency, and circulation signature markers in different subsets of T cells.

**A**

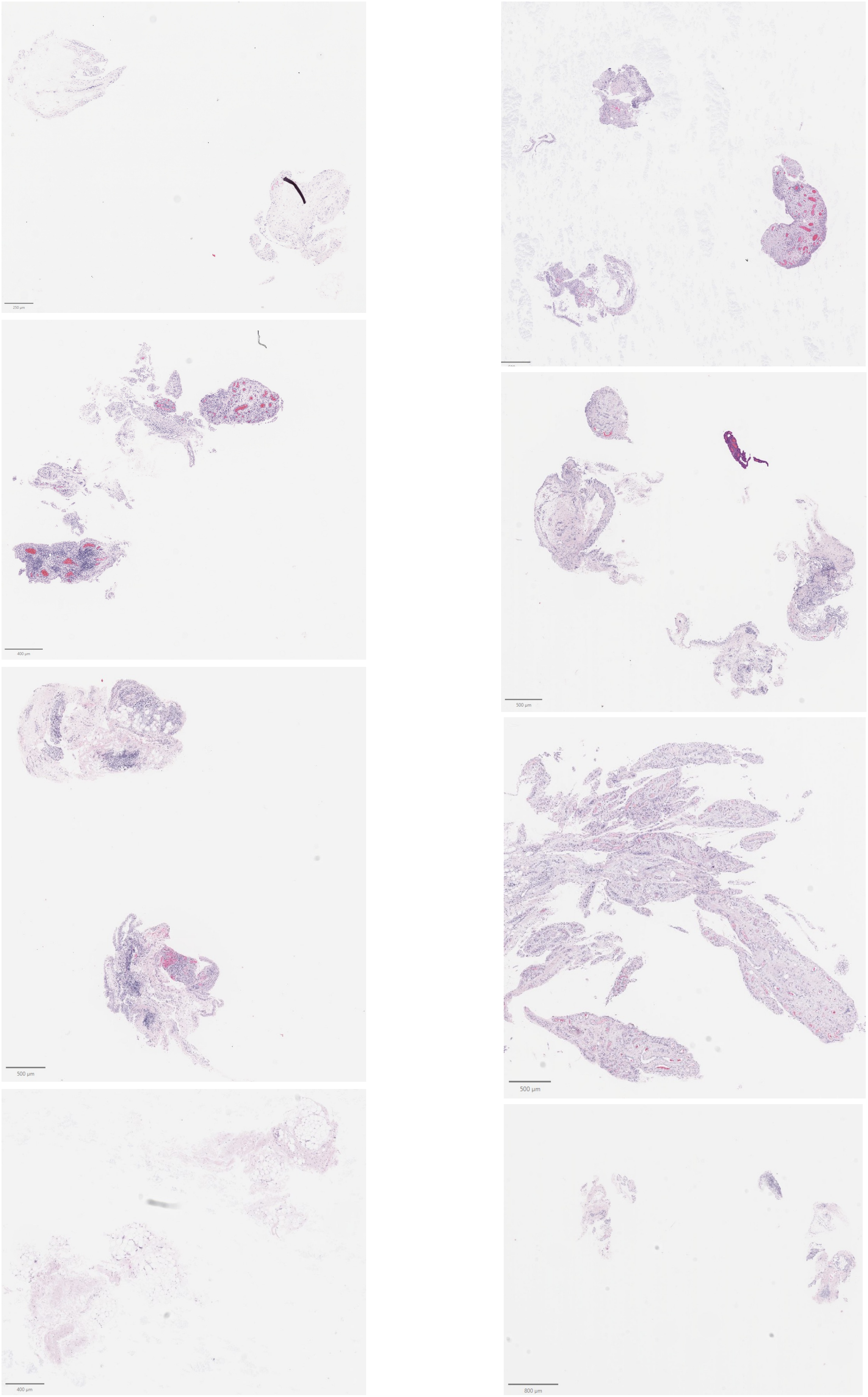


**B**

Please find 80 images attached in the separate PDF file named Figure S8B.

**Figure S8. Hematoxylin and eosin stain and images for the CosMx spatial profiling.** (A) H&E stain for eight tissues from 4 PsA patients. (B) 80 fields of view (FoVs) images scanned.

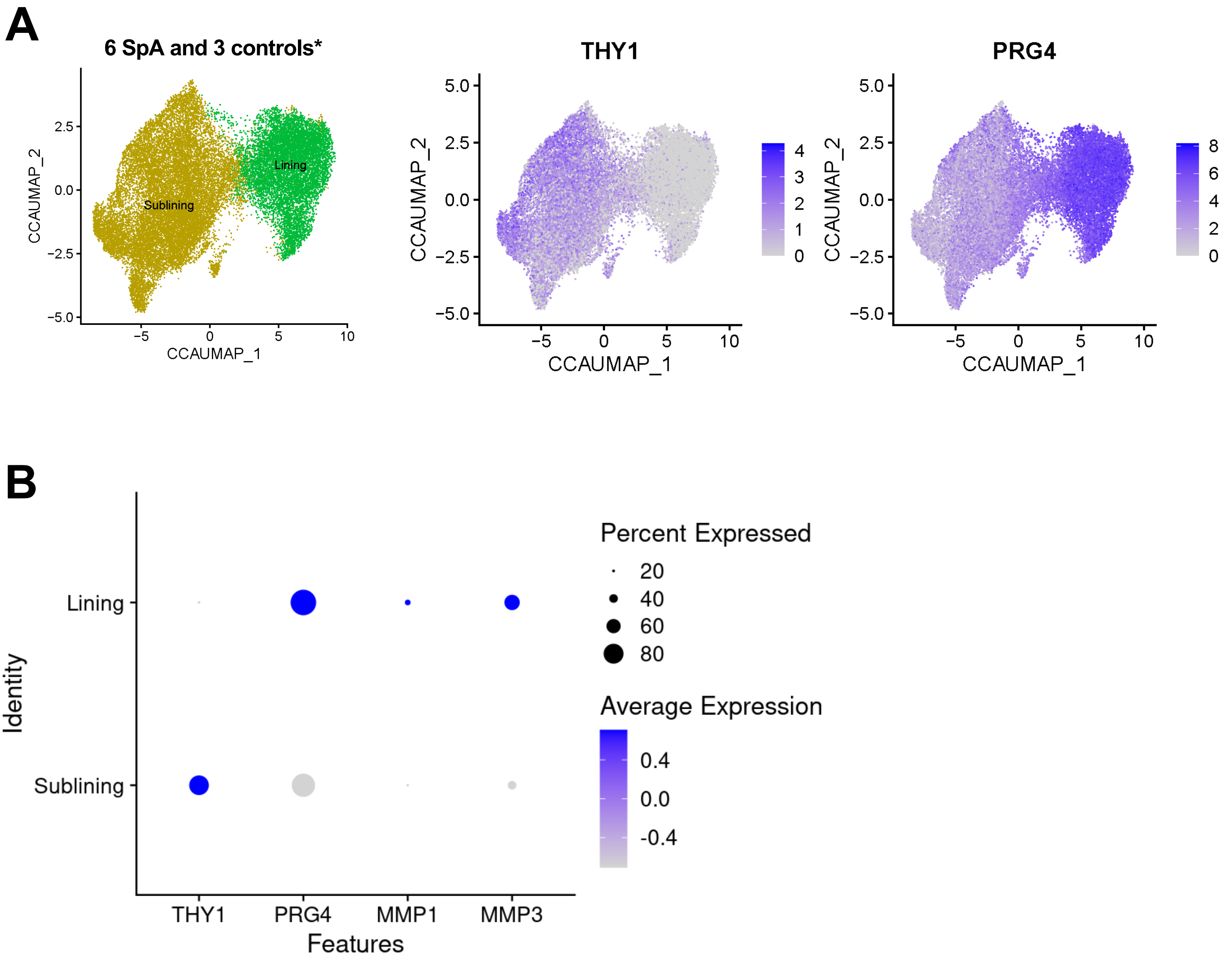


**Figure S9.** ***THY1-PRG4+*** **lining and** ***THY1+PRG4-*** **sublining fibroblasts represent two major subsets of synovial fibroblasts.**

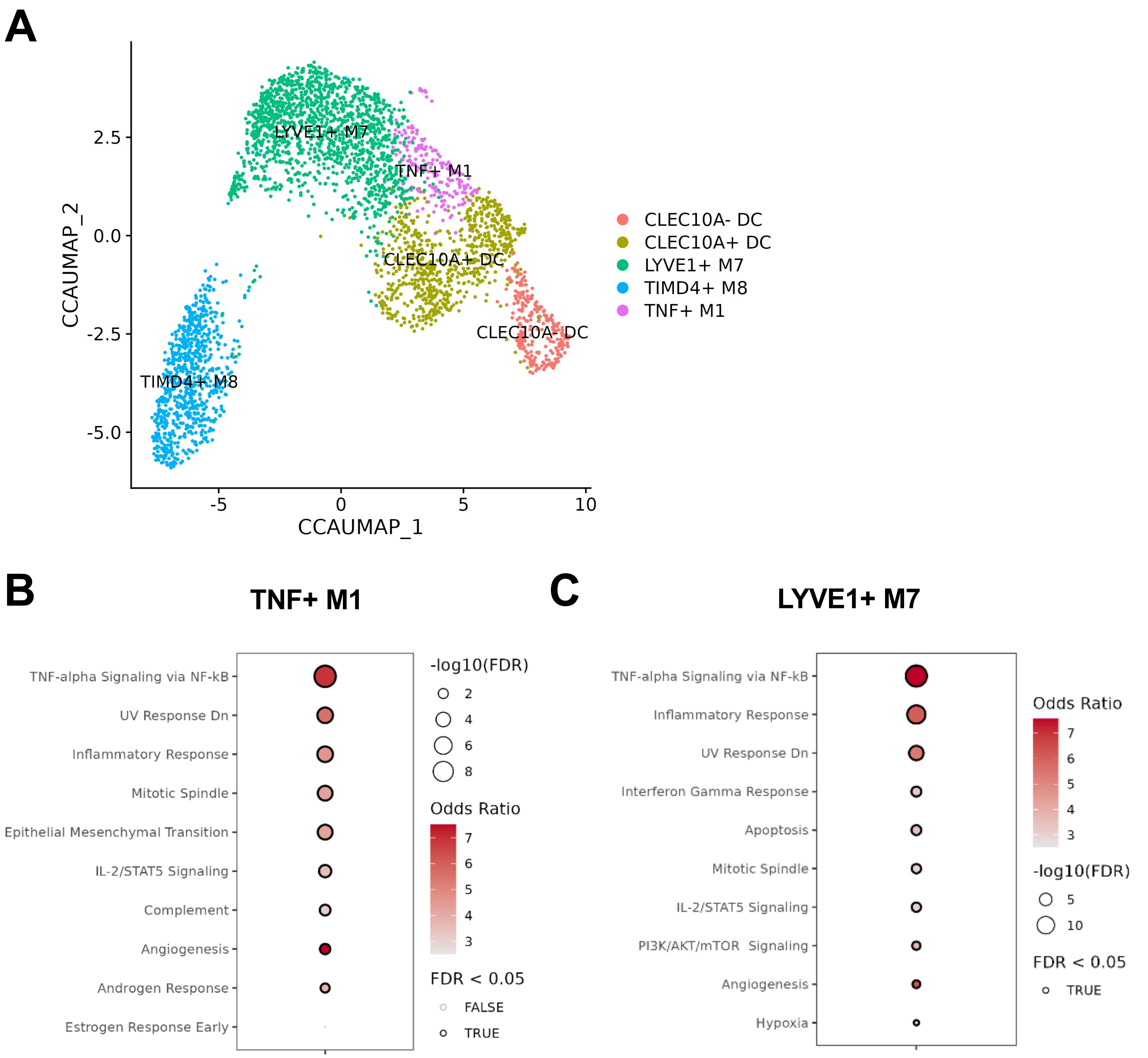


**Figure S10. Proinflammatory pathways are enriched in TNF+ M1 and LYVE1+ M7 cell populations from SpA.** (A) UMAP visualization of subsets of myeloid cells from healthy synovial tissues. (B) and (C) show the enriched pathways in TNF+ M1 and LYVE1+ M7 respectively. All up-regulated genes were used for enrichment analysis using MSigDB database.

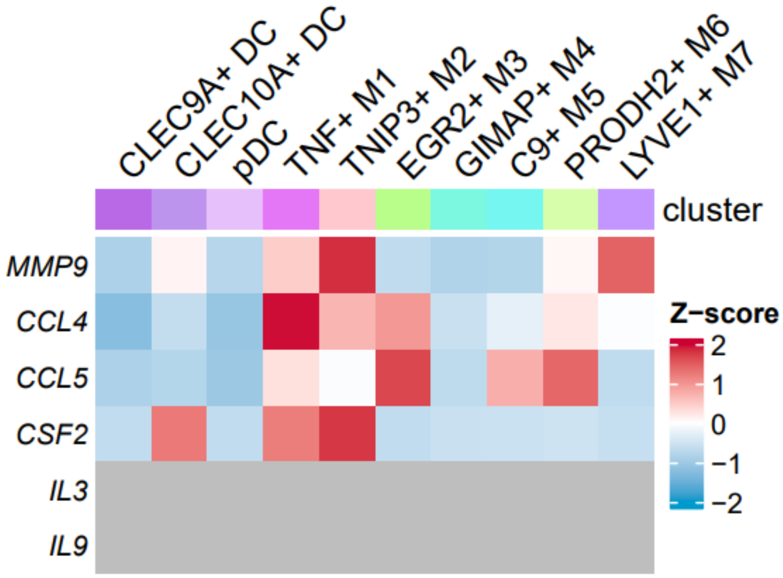


**Figure S11. Expression of IL-17-induced genes in different myeloid subsets.** Heatmap of normalized and scaled expression of previously reported IL-17-induced genes in different subsets of myeloid cells.

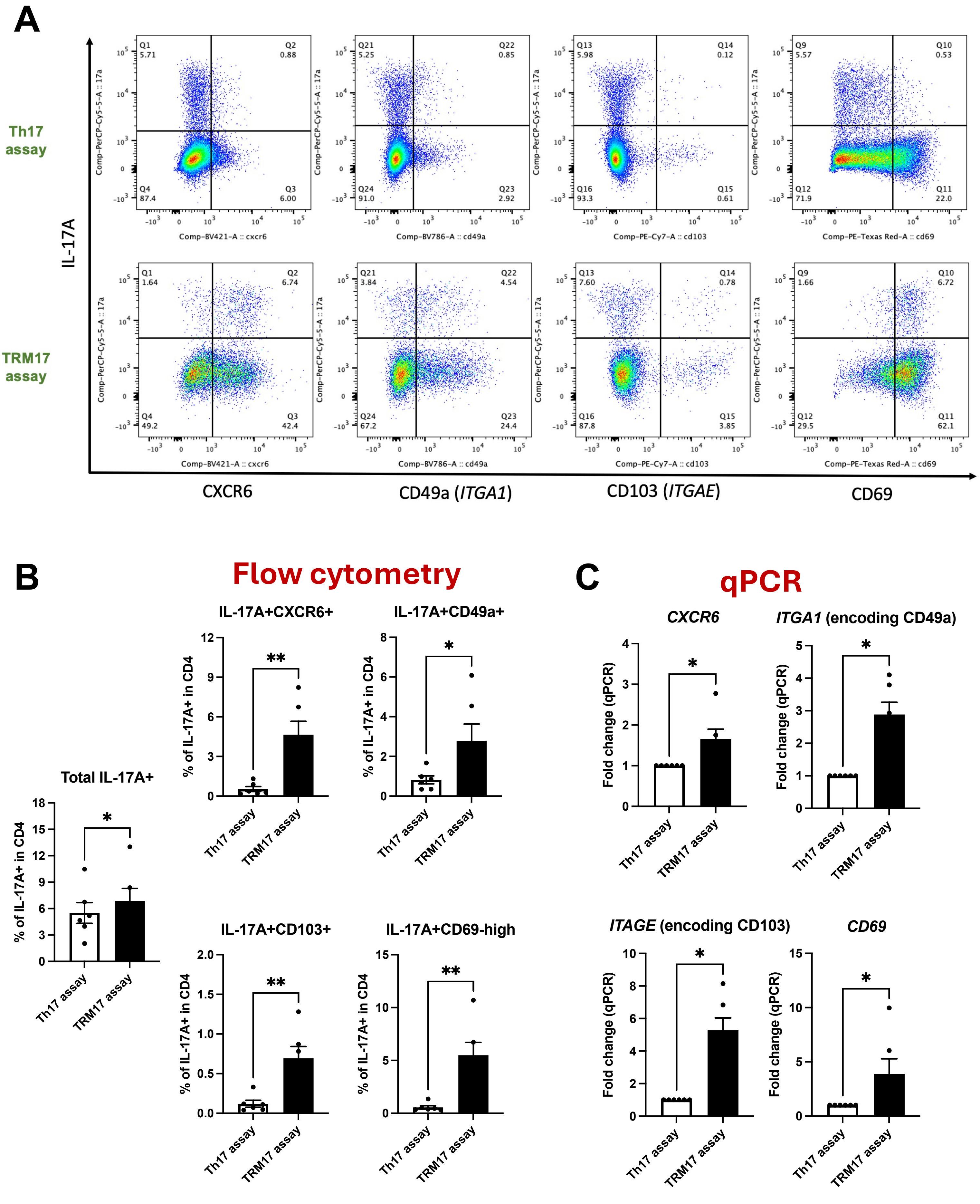


**Figure S12. The TRM17 assay induces higher expression of TRM markers in Th17 cells compared to the Th17 assay.**

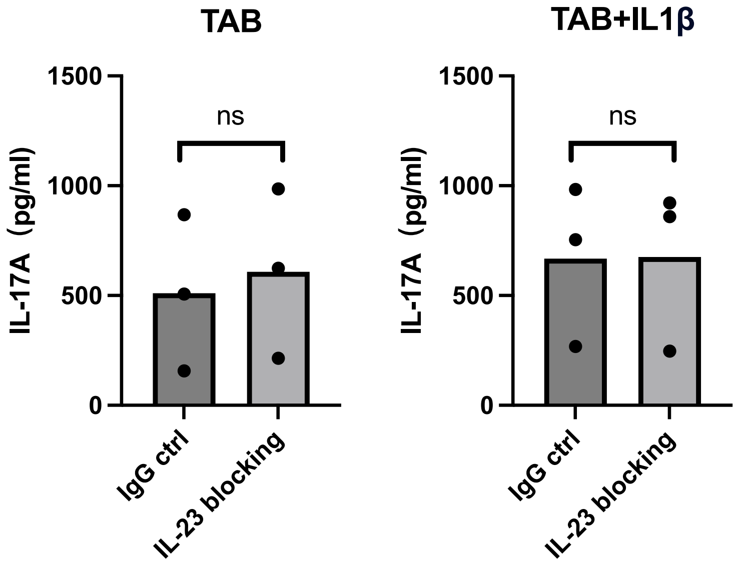


**Figure S13. IL-23 blockade does not reduce the secretion of IL-17A by TRM17-like cells.** The levels of cytokines IL-17A produced by TRM17-like cells treated with 10 µg/ml IL-23 blocking antibody or IgG control in the presence of TAB or TAB and IL-1β. The P-value was assessed using the paired two-tailed Student's t-test (* $P \leq 0.05$).

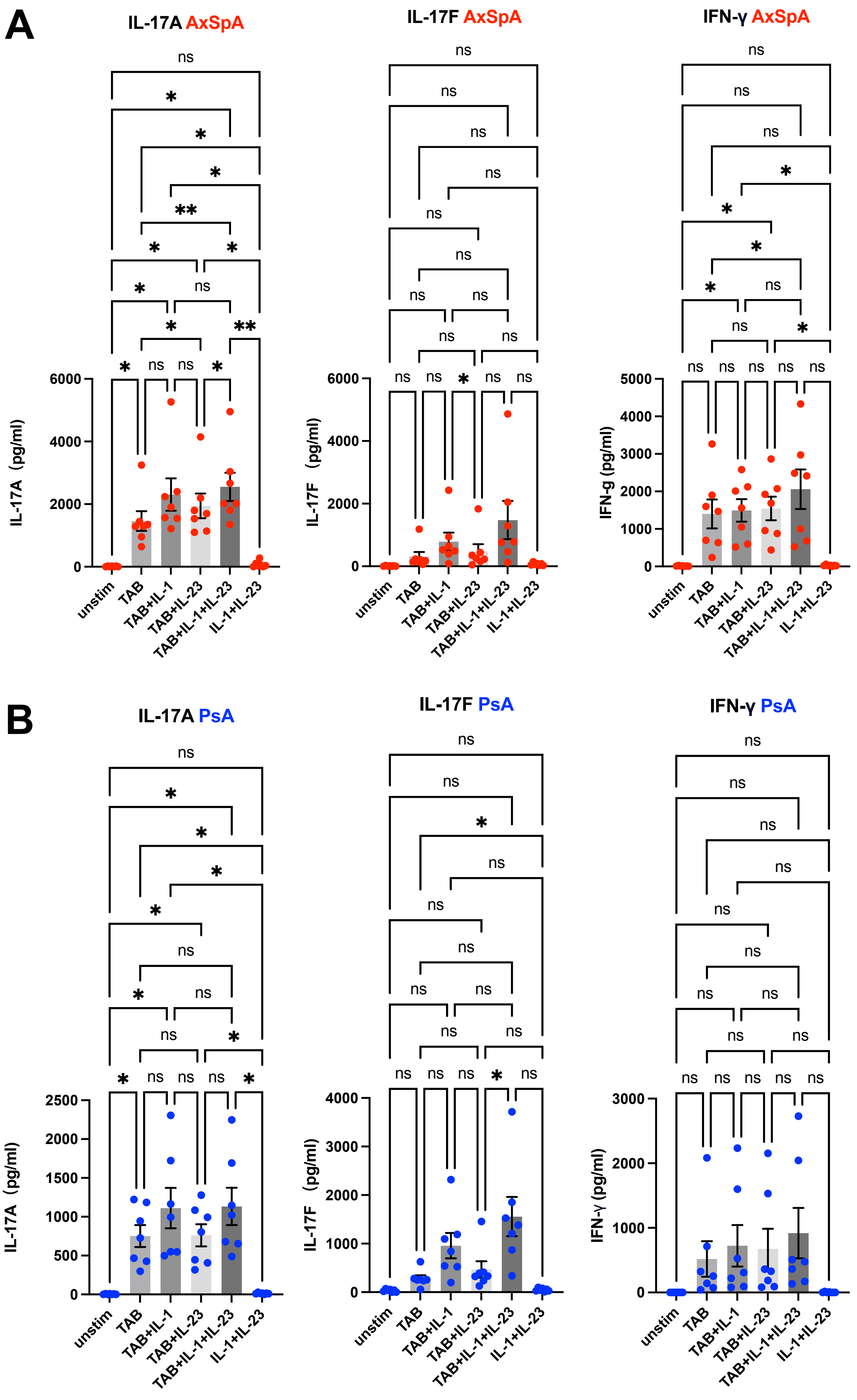


**Figure S14. T-cell receptor engagement and cytokine stimulations co-drive the production of IL-17A/F by in vitro generated TRM17 - like cells.** The secretion of cytokines by TRM17-like cells stimulated with vehicle control, cytokines (IL-1β and IL-23), T-cell activation beads (TAB) or the combinations of cytokine and TAB. Data from 7 AxSpA (A) and 7 PsA (B) patients are shown. Statistical significance was assessed using one-way ANOVA (* $P \leq 0.05$, ** $P \leq 0.01$).

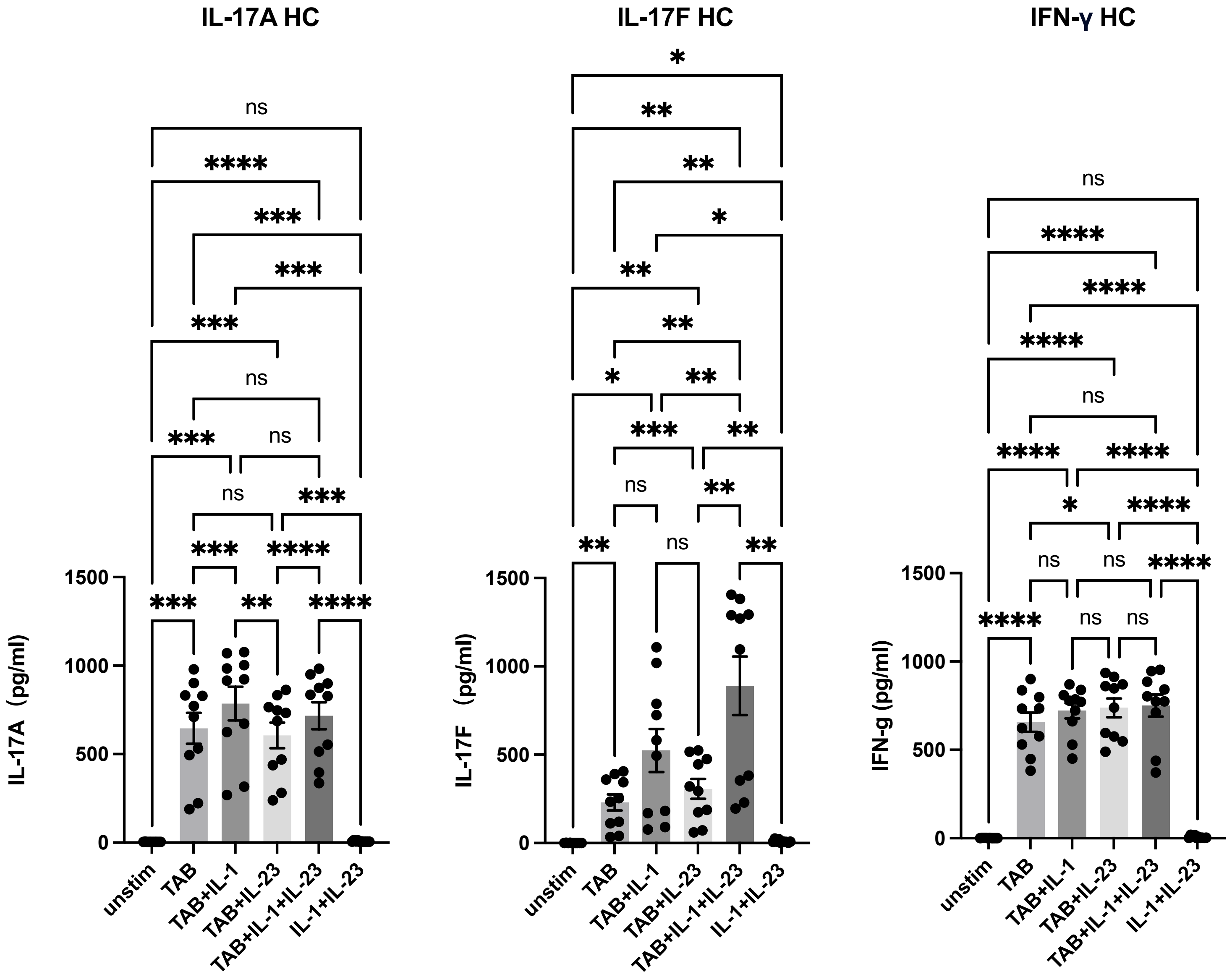


**Figure S15. T-cell receptor engagement and cytokine stimulations co-drive the production of IL-17A/F by in vitro generated TRM17 - like cells from healthy controls.** Statistical significance was assessed using one-way ANOVA (* $P \leq 0.05$, ** $P \leq 0.01$, *** $P \leq 0.001$, **** $P \leq 0.0001$).

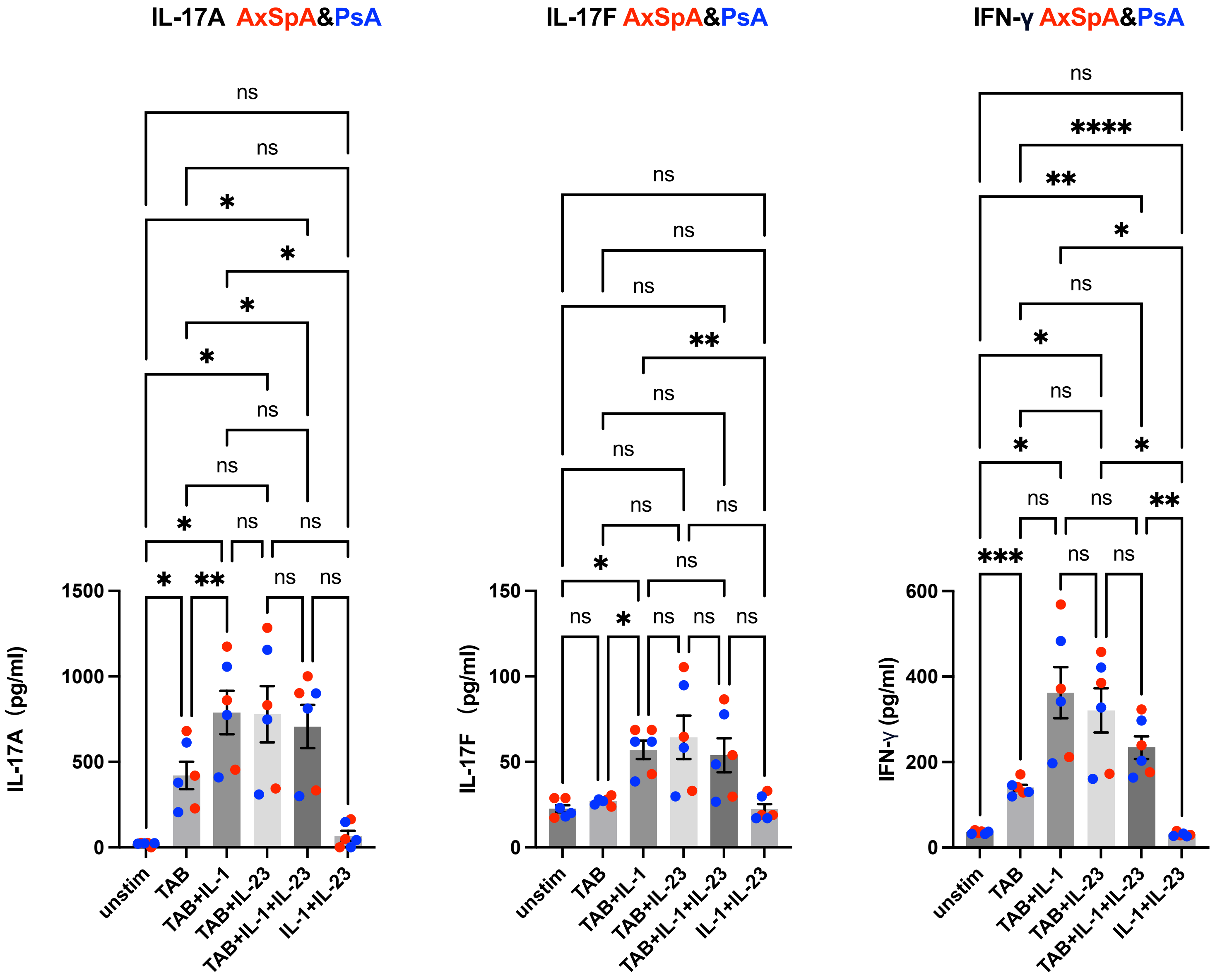


**Figure S16. Response of cells from Th17 assay to stimulations.** Statistical significance was assessed using one-way ANOVA (* $P \leq 0.05$, ** $P \leq 0.01$, *** $P \leq 0.001$, **** $P \leq 0.0001$).

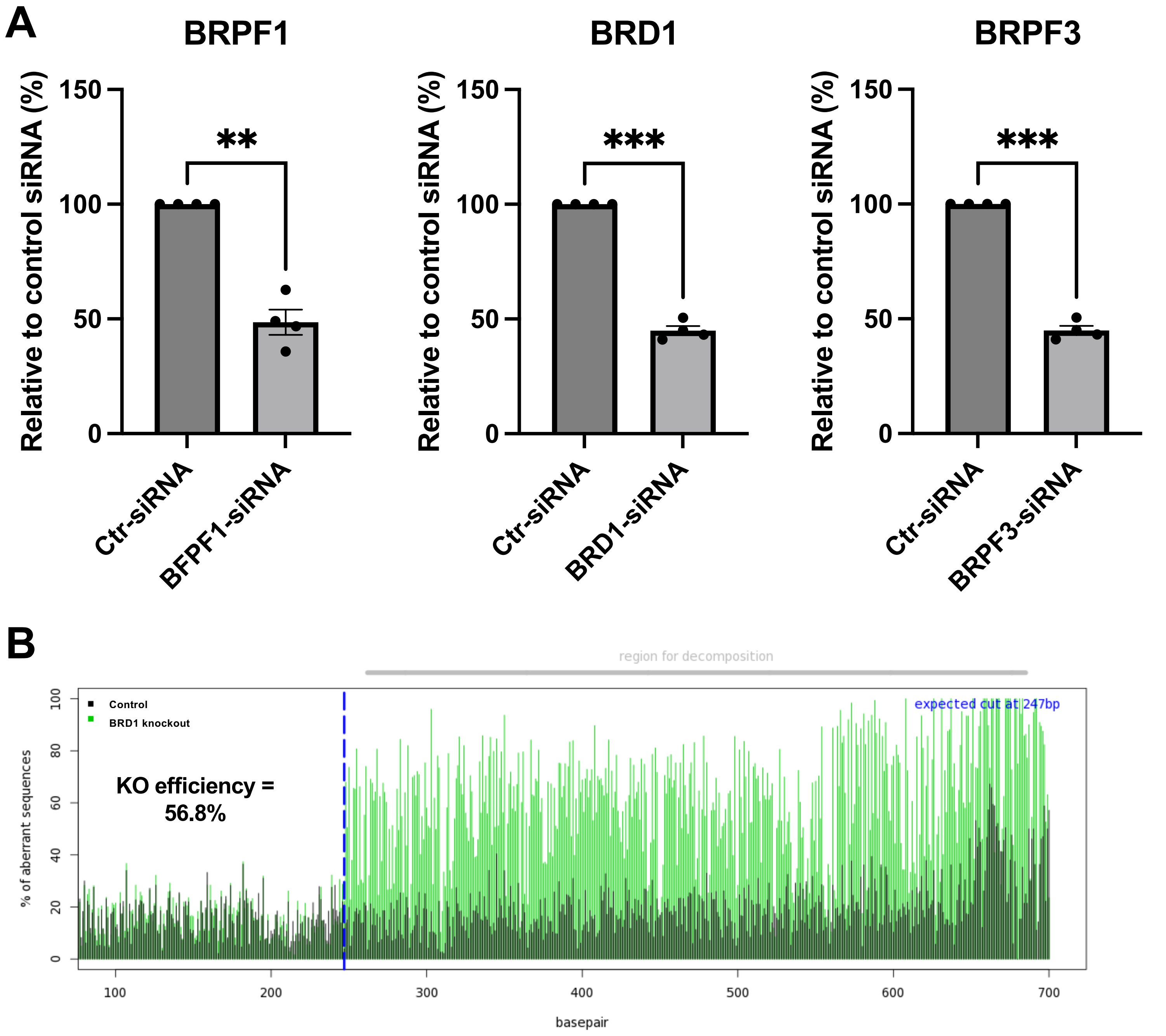


**Figure S17. siRNA Gene Silencing and CRISPR Knockout Efficiency.** (A) Efficiency of BRPF1, BRD1, and BRPF3 siRNA silencing was measured using qPCR. (B) Efficiency of BRD1 knockout was assessed using the classical TIDE tool (http://shinyapps.datacurators.nl/tide/). Specifically, genomic DNA from the target region was PCR-amplified, followed by sequencing. The Knockout efficiency was calculated by comparing the BRD1 knockout sample to control unedited cells using TIDE. For (A) Statistical significance was determined using the paired two-tailed Student's t-test (**$P \leq 0.01$, ***$P \leq 0.001$). Means +/- SEM are shown.

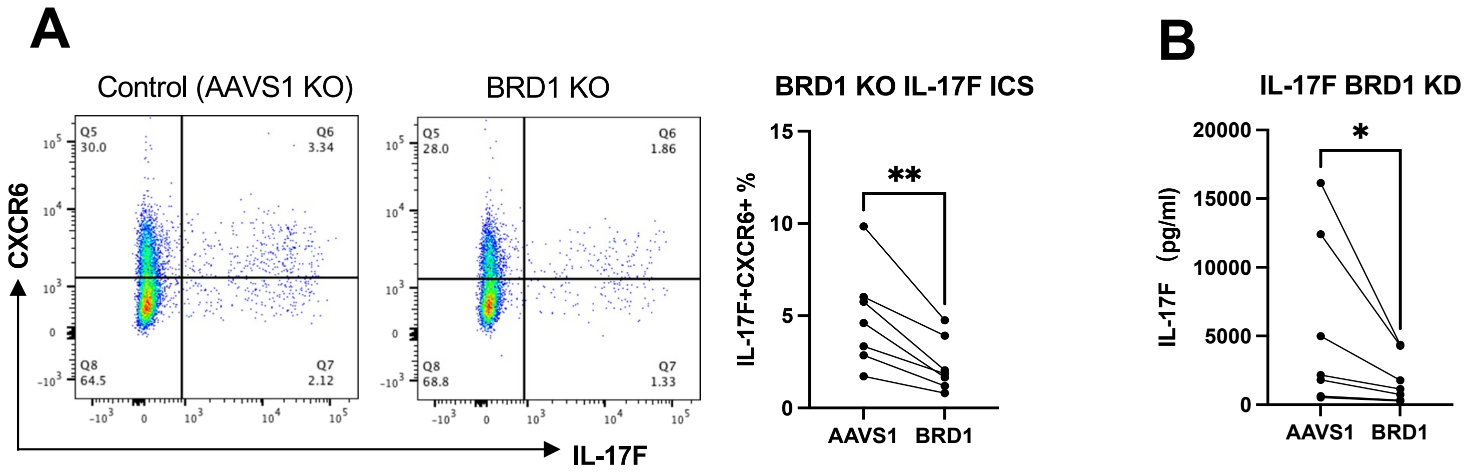


**Figure S18. Inhibition of IL-17F+CXCR6+ cell frequency and IL-17F production in TRM17-like cells by BRD1 KO.** Effect of BRD1 knockout on the generation of TRM17-like cells and their IL-17F production upon antigen re-call (data from 7 SpA patients). The changes of Th17 frequency (based on IL-17F intracellular cytokine staining) are shown (A) The P-value was assessed using the paired two-tailed Student's t-test (* P≤ 0.05; ** P≤ 0.01). The level of IL-17F secretion (ELISA) relative to vehicle control (DMSO) is shown (B). The P-value was assessed using the Wilcoxon test (* P≤ 0.05).
